# SVEP1 enables efficient binding of Angiopoietin-2 to the TIE1 receptor, allowing receptor phosphorylation and downstream signaling

**DOI:** 10.1101/2025.03.17.643633

**Authors:** Katharina Uphoff, Melina Hußmann, Oliver Peters, Matthias Mörgelin, Fabian Metzen, Wolfgang M.J. Obermann, J. Fernando Bazan, Jörg Stetefeld, Manuel Koch, Stefan Schulte-Merker, Dörte Schulte-Ostermann

## Abstract

The molecular mechanisms that drive (lymph-)angiogenesis are crucial to understand diseases, such as lymphedema, that are caused due to malformations of the lymphatic vasculature. Recently, an interaction between the secreted protein Svep1, a key regulator in lymphangiogenesis, and the transmembrane receptor Tie1 was shown in zebrafish, human, and mice. Here, guided by *in silico* AlphaFold-multimer structure predictions of SVEP1 complexes, we assert with protein binding studies that the human CCP20 domain is the primary binding site for TIE1. We further demonstrate that SVEP1 mediates strong binding of ANG2 and TIE1, and that combined stimulation of hdLECs with SVEP1 and ANG2 leads to phosphorylation of TIE1. TIE1 activation by SVEP1 and ANG2 enables downstream signaling and, in turn, potentiates nuclear exclusion of FOXO1 and phosphorylation of AKT compared to SVEP1 or ANG2 alone. We present a model in which ANG1/2 dimers bind to both SVEP1 and TIE1, resulting in the recruitment of multiple TIE1 receptor molecules to a multimeric complex at the cell membrane, potentially amplifying its signaling capacity.

## Introduction

Lymphatic vessels are closely associated with blood vessels, and malformations of the lymphatic vasculature lead to lymphedema, obesity, or chronic inflammatory diseases (Mäkinen et al. 2021; Petrova and Koh 2018; Sunil Kumar et al. 2025). Next to the well-described lymphatic VEGFC/VEGFR3 pathway (Hogan et al. 2009; Bos et al. 2011; Le Guen et al. 2014; Jeltsch et al. 2014; Roukens et al. 2015), it was shown that Sushi, von Willebrand factor type A, EGF, and pentraxin domain containing protein 1 (SVEP1), also referred to as Polydom, is another key player in lymphangiogenesis (Karpanen et al. 2017; Morooka et al. 2017). Svep1 encodes a 3571 amino acid long extracellular matrix protein of different domains like von Willebrand factor type A domain (vWF), ephrin-receptor like domains, complex control protein (CCP) domains, and Hyalin repeats at the N-terminus. The C-terminus mainly consists of CCP and EGF domains. Svep1 is expressed in mesenchymal cells and functions non-cell-autonomously (Karpanen et al. 2017; Morooka et al. 2017). Only recently, it was discovered that SVEP1 binds to TIE1 (Hußmann et al. 2023; Sato-Nishiuchi et al. 2023), and we were able to provide *in vivo* evidence for the genetic interaction of *svep1* and *tie1* in zebrafish. Multiple phenotypic hallmarks of *svep1* mutants are phenocopied by *tie1*, but not *tie2* mutants, including (among others) a reduced number of parachordal lymphangioblasts (PLs) (Hußmann et al. 2023). Furthermore, a specific aspect of the facial lymphatics in zebrafish, the facial collecting lymphatic vessel (FCLV), develops in a *svep1* and *tie1* dependent, but *vegfc/vegfr3* and *tie2* independent manner (Hußmann et al. 2023).

The discovery of the novel molecular interaction between SVEP1 and TIE1 provided an invitation to further investigate how SVEP1 is linked to the already well-known ANG/TIE pathway. The ANG/TIE pathway triggers a signaling cascade, which is required for lymphatic and blood vessel development, and has a unique role in maintaining vascular stability. Two receptor tyrosine kinases (RTKs), tyrosine kinase with Ig and EGF domains (TIE1) and tyrosine endothelial kinase (TEK), also known as TIE2 (Dumont et al. 1993; Partanen et al. 1992), are activated through binding of angiopoietin ligands, including ANG1 and ANG2 (Davis et al. 1996; Maisonpierre et al. 1997). TIE1 and TIE2 exhibit a high degree of homology with a globular head domain consisting of three immunoglobulin-like (Ig) domains and three epidermal growth factor-like (EGF) modules and a short stalk formed by three fibronectin type III repeats, while the N-terminal two Ig domains of TIE2 harbor the angiopoietin binding site (Macdonald et al. 2006). TIE2 is considered the main player in angiogenesis, and its phosphorylation, triggered by ANG1 binding, induces downstream signaling via PI3K/Akt, which leads to inhibition of the transcription factor Forkhead box protein O1 (FOXO1) and repression of FOXO1 target genes -such as *Ang2* (Kim et al. 2000, Daly et al. 2004). Although TIE2 is needed for lymphangiogenesis in mouse embryos, with genetic deletions of *Tie2* in lymphatic vessels leading to subcutaneous edema (Souma et al. 2018), postnatal knock-out of *Tie2* in lymphatics of newborn mice only has an effect on lymphatic collecting vessels but not on cutaneous lymphatic capillaries (Korhonen et al. 2022; Shen et al. 2014). While ANG1 is an agonist of TIE2, ANG2 acts as a context-dependent weak agonist or antagonist for TIE2 that can inhibit the ANG1–TIE2 signaling axis (Saharinen, Eklund, and Alitalo 2017). TIE1 blocks the signaling cascade in a context dependent manner by forming heterodimers with TIE2 (Hansen et al. 2010; Marron et al. 2000; Saharinen et al. 2005; Savant et al. 2015; Seegar et al. 2010). On the other hand, *Tie1* deletion leads to defects in the lymphatic capillary network and primary lymphatic network remodeling (Korhonen et al. 2022; Shen et al. 2014).

In the present study we extend our understanding of SVEP1 binding to TIE1, utilizing *in silico*, *in vitro*, and *in vivo* techniques. We show that the binding of ANG2 to TIE1 is enhanced by SVEP1 leading to phosphorylation and downstream signaling of TIE1, thereby potentiating phosphorylation of AKT and nuclear exclusion of FOXO1. Our study helps to understand the interaction of important signaling pathway members involved in (lymph-)angiogenesis and provides a model that suggests local TIE1 multimerization through bridging functions of ANG1/2 in the presence of SVEP1.

## Results

### SVEP1 binds to the D1-D2 domain of TIE1

Previously, it had been shown that solid-phase binding assays of recombinant full-length murine SVEP1 protein display a significant binding of SVEP1 to TIE1, but not TIE2 (Sato-Nishiuchi et al. 2023). In a binary setup, i.e., without other proteins being present, we confirmed this observation utilizing surface plasmon resonance (SPR) technology and negative staining assays (Figure 1 C and D). Furthermore, and since AlphaFold2 (AF2) and AF3 are remarkably accurate predictors of evolutionarily conserved protein-protein interactions, we used their multimer capabilities to search for significant contact points between the full-length, 3571 amino acid human SVEP1 chain and potential binding partners TIE1 and TIE2. As shown by the comparative PAE (Predicted Aligned Error, a prediction quality and model confidence metric) plots of the full-length SVEP1 matches with TIE1 and TIE2 (Figure 1A, B), we observe clustering between the headpiece or N-terminal D1-D4 domains of TIE1 and the CCP19-21 stretch of SVEP1, in line with the results of Sato-Nishiuchi et al. (2023) which center on the CCP20 domain of murine SVEP1 for TIE1 capture. D1 and D2 refer to the two N-terminal Ig domains, D3 refers to the three EGF domains, and D4 to the third Ig domain of TIE1 or TIE2. By contrast, the PAE plot for the D1-D4 segment of TIE2 fails to show an equivalent clustering with the CCP19-21 stretch of full-length SVEP1. The more focused interrogation and modeling of equivalent N-terminal TIE1 and TIE2 ecto-domains with the isolated CCP19-CCP21 domain fragment of SVEP1 (using AF2.3 multimer) affirms the interaction interface of SVEP1 to TIE1, positioning the groove between the side-by-side packed Ig-like D1-D2 domains of TIE1 in close contact with the CCP20 domain of SVEP1, revealing an interaction surface area of 851,5 Å^2^ marked by 8 H-bonds and 2 salt bridges by PDBePISA analysis (Supplementary Figure 1.1). The SVEP1-binding domains of TIE1 are buttressed by a Cys-rich D3 domain that links to an outstretched D4 Ig module that further connects to a linear array of D5-D7 FnIII domains N-terminal to the hydrophobic TM helix. Key regions and contacts illustrating salt bridges, hydrogen bond network surface, electrostatic potential representation and hydrophobicity surface representations are shown in supplementary Figure 1.2. In this closer examination of molecular fragments of TIE2 and SVEP1, AF2.3 multimer scores the interface lower in both ipTM (interface predicted template modelling) and actifpTM (actual interface predicted template modelling) metrics (Supplementary Figure 1.1) and qualitatively weaker binding to the CCP19-21 fragment of SVEP1. The resulting SVEP1-TIE2 model shows the D1 and D2 domains of TIE2 perched between the CCP20 and CCP21 domains of SVEP1 (Figure 1B). While PDBePISA analysis shows a similar interface surface area of 889.4 Å^2^, only 2 H-bonds and 1 salt bridges are observed, suggesting a weaker interface. To verify the modelling results, we performed SPR assays with a 70 kDa version of the SVEP1 protein that contains the CCP20 domain (Figure 1C). We observed interaction of SVEP1 with TIE1, but not with TIE2 in the absence of other proteins. Furthermore, utilizing negative staining and transmission electron microscopy, we were able to directly visualize the attachment of SVEP1 and TIE1 (Figure 1D). As shown in supplementary Figure 1.2, modelling of SVEP1 and TIE1 suggested P202 and L203 of TIE1 as a key contact region. Accordingly, a TIE1 (P202L-L203F) mutant was not able to bind to SVEP1 (Supplementary Figure 1.3B). In addition and in agreement with a previous study, a SVEP1 (E2568A-G2569A) mutant in which two crucial amino acids in the CCP20 domain are exchanged could not bind to TIE1 (Supplementary Figure 1.3A).

**Figure 1:**
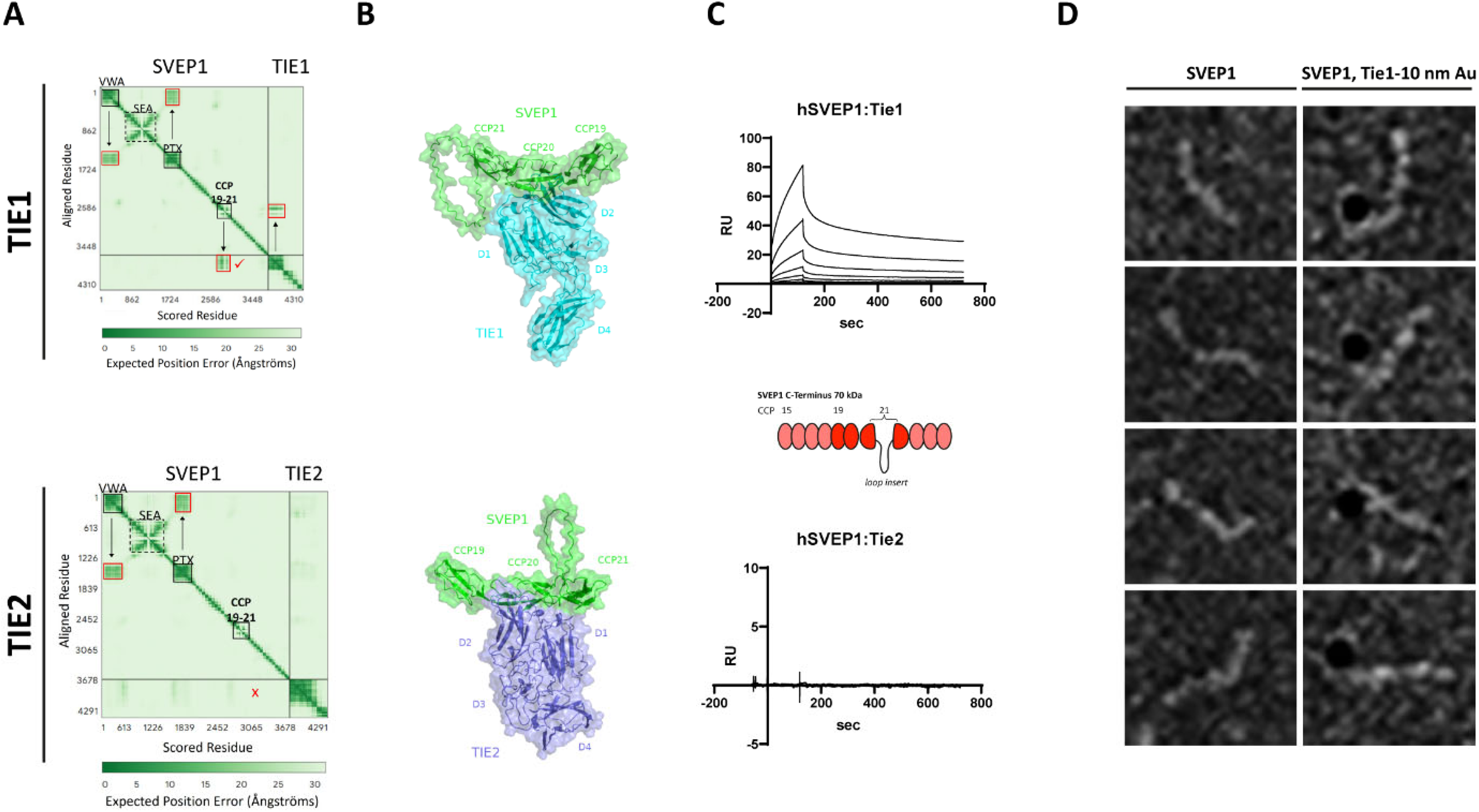
The CCP20 domain of SVEP1 contacts the D1-D2 headpiece domains of TIE1. (**A**) Comparative PAE plots of full-length SVEP1 and the TIE1 ectodomain displays a primary binding site for TIE1 (domains D1-D2) by AF3-multimer modeling, that localizes to the CCP19-21 stretch, notably encompassing CCP20.. The PAE plot for full-length SVEP1 and the TIE2 ectodomain does not show an equivalent prediction for the same CCP19-21 region. Labelled SVEP1 domains: PTX (Pentraxin), SEA (sperm protein, enterokinase and agrin) module, VWA (von Willebrand factor type A), statistics flSVEP1-ectoTIE1 pLDDT = 66.1; pTM = 0.34; ipTM = 0.48; statistics flSVEP1-ectoTIE2 pLDDT = 0.68, ipTM = 0.28, pTM = 0.34 (**B**) A discrete AF2.3 model of TIE1 bound to CCP19-21 shows that the Ig-like D1-D2 headpiece domains of TIE1 (linked by a Cys-rich D3 to an ‘open’ or flared-out D4 Ig module) are in contact with CCP20. By contrast, the AF2.3 model of TIE2’s N-terminal domains is more loosely perched between the CCP20 and CCP21 domains with a qualitatively lower affinity due to reduced sidechain contacts. (**C**) Quantitative analysis of TIE1-SVEP1 interaction using SPR assays. In the absence of other proteins, a 70kD version of SVEP1 (CCP15-CCP24, amino acids: 2261-2890; depicted in diagram fashion) containing the CCP20 domain binds TIE1 (K_D_ = 62.5 nM), but does not bind TIE2. (**D**) Negative staining shows binding of human C-term SVEP1-Strep II (reaching from CCP15 to the C-Term) and TIE1, labelled with 10nm Au. PAE: Predicted aligned error; AF3: AlphaFold 3; CCP: complement control protein (sushi repeats).

### ANG1/2 strengthens the binding capacity of SVEP1 and TIE1

Murine SVEP1 binds ANG1 and ANG2 *in vitro* (Morooka et al. 2017), and it has long been known that ANG1 and ANG2 bind to TIE2 (Davis et al. 1996; Maisonpierre et al. 1997). Both AF2 and AF3 multimer models effectively capture TIE1 in a complex with SVEP1 CCP20, so we next explored how the addition of ANG1/2 would affect this assembly (Figure 2A). The affinity of TIE1 for SVEP1 is significantly increased by the simultaneous binding of the C-terminal domains of ANG1 or ANG2 to the CCP20 module, as suggested by the increased PAE and ipTM scores, which improve on the binary coupling of just TIE1 to SVEP1 (Supplementary Figure 2.1). By contrast (not shown), TIE2 was not predicted to stably co-bind with ANG1/2 to CCP19-CCP21 of SVEP1 because the weaker binding of TIE2 with CCP19-CCP21 of SVEP1 fails to nucleate the formation of a ternary complex. The conformation of the TIE1-ANG1/2 heteromers in the SVEP1-bound pose resembles the solved X-ray structures of TIE2 bound to either ANG1 or ANG2 (PDB codes 4K0V and 2GY7, respectively, in the absence of any experimental TIE1-ANG complexes) (Supplementary Figure 2.1), and furthermore, the equivalent ANG1/2-binding epitopes in TIE1 are not blocked by SVEP1 binding. Whereas the two TIE2 X-ray complexes show a buried surface area of ∼1200 Å^2^ (with 2 H-bonds and 2 salt bridges) against ANG1/2, the addition of the SVEP1-TIE1 contact produces a combined ∼1800 Å^2^ interaction surface area as well as new contacts between SVEP1-ANG1 and SVEP1-ANG2 (3 H-bonds and 1 H-bond, respectively) (Supplementary Figure 2.1). For this reason, SVEP1’s CCP20 module acts as a binding partner for TIE1, but not TIE2, when interacting with ANG1/2.

**Figure 2:**
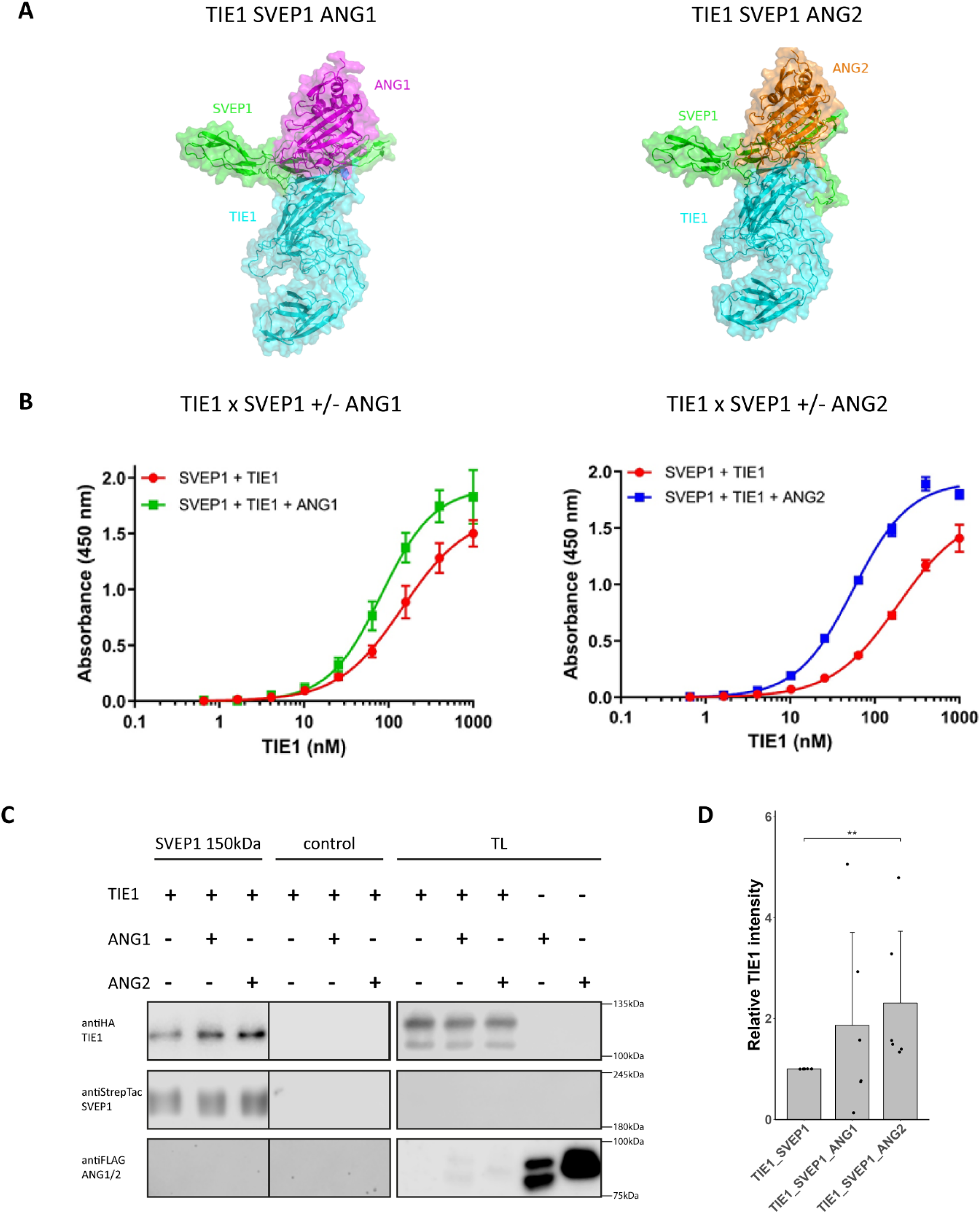
ANG1/2 strengthens the binding capacity of SVEP1 and TIE1. (**A**) AF2.3-modeling of the SVEP1 CCP19-21 domain and TIE1 together with ANG1/2. The affinity of TIE1 for SVEP1 is considerably heightened by the co-binding of ANG1 or ANG2 C-terminal domains to the CCP20 module. (**B**) The Interaction between the complete C-terminal SVEP1 (spanning from the EGF1 domain to C-terminus) and the ectodomain of TIE1 was analyzed by ELISA binding assays. SVEP1 proteins were immobilized on 96 well plastic plates and overlayed with serial dilutions of TIE1 fused to an ALFA-tag, in the absence or presence of equal amounts of ANG2 as indicated. Bound TIE1 proteins were detected by HRP conjugated anti-ALFA-tag nanobody. ANG2 binding was detected using an HRP conjugated anti-Biotin antibody against the biotinylated protein. KD values are lower in the presence of ANG1 (K_D_=82,4) or ANG2 (K_D_=55,6), reflecting a higher affinity than SVEP1 and TIE1 in the absence of angiopoietins (K_D_=189,0). (**C**) Co-immunoprecipitation of human TIE1 co-transfected with either human ANG1 or ANG2 in 293T HEK cells. Human SVEP1-Strep II (a 150 kDa version reaching from CCP15 to the C-Term) was immunoprecipitated, and associated TIE1 and ANG1/2 were detected via Western blot analysis. (**D**) Quantification of the TIE1 levels with SVEP1 levels as a reference confirmed a significant increase in binding affinity of SVEP1 and TIE1 together with ANG2 (**p.adj = 0.008; Mann–Whitney U test; Values are presented as means ± SD; individual data points for each experiment). TL, total lysate.

ELISA assays verified the model prediction as the presence of either ANG1 or ANG2 led to a lower KD value (SVEP1, TIE1 (KD = 146,9) vs. SVEP1, TIE1, ANG1 (KD = 82,4); SVEP1, TIE1 (KD = 189) vs. SVEP1, TIE1, ANG2 (KD = 55,6)), reflecting a higher affinity (Figure 2B). Furthermore, co-transfection of either ANG1 or ANG2 together with TIE1 and subsequent immunoprecipitation of SVEP1 bound to beads verified the model prediction (Figure 2C). Whereas SVEP1 levels in all samples were similar, the level of TIE1 bound to SVEP1 significantly increased when ANG1/2 was present in the lysate. TIE1 co-transfected with ANG1 led to a slight, but not significant, increase in TIE1 levels. However, after transfection of TIE1, ANG1 and ANG2 levels are decreased in the total lysate. Thus, we were not able to detect ANG1 or ANG2 after immunoprecipitation.

Furthermore, negative staining shows binding of TIE1 and ANG1/2 to SVEP1 (Supplementary figure2.2)

### Stimulation by SVEP1 and ANG1/ANG2 leads to phosphorylation of TIE1, nuclear exclusion of FOXO1, and phosphorylation of AKT

In previous studies, phosphorylation of TIE1 could not be detected after stimulation of cells with SVEP1 (Sato-Nishiuchi et al. 2023) or ANG2 (Saharinen et al. 2005) only, whereas stimulation of EA.hy926 immortalized hybrid HUVECs with ANG1, ANG4, and COMP-ANG1 leads to phosphorylation of TIE1 (Saharinen et al. 2005). Our aim was to test whether cooperative binding of SVEP1 and angiopoietins to TIE1 can lead to TIE1 phosphorylation, where SVEP1 constitutes the connecting link between TIE1 and ANG2. Indeed, after stimulation of hdLECs with both SVEP1 and ANG2, robust phosphorylation of TIE1 at residue pY^1007^ was detected (Figure 3A and D). Also, stimulation with SVEP1 alone has a slight effect on the phosphorylation of TIE1, possibly because hdLECs express ANG2 (Korhonen et al. 2022). To further test whether SVEP1-ANG2 stimulation leads to downstream signaling, we analyzed pAKT levels. Stimulation with either ANG2 or SVEP1 led to phosphorylation of AKT, which was amplified by simultaneous stimulation with ANG2 and SVEP1. The effect of SVEP1 stimulation on AKT signaling could be blocked by an ANG2 blocking antibody, suggesting that SVEP1 does not induce signaling on its own, but enables signaling of ANG2 (Figure 3B and E). Knock-down of either TIE1 or TIE2 could reduce activation of pAKT, leading to the assumption that both TIE1 and TIE2 are necessary for the phosphorylation of AKT after SVEP1-ANG2 stimulation and thus downstream signaling. Furthermore, we analyzed FOXO1 nuclear exclusion, a downstream event of the PI3K/AKT pathway, by stimulating hdLECs with either ANG2 or SVEP1, or both proteins in combination, and testing for nuclear versus cytoplasmic FOXO1 ratios. It has already been shown that SVEP1 activates the PI3K-Akt signaling pathway *in vitro*, with nuclear exclusion of FOXO1 (Sato-Nishiuchi et al. 2023). We observed enhanced nuclear exclusion of FOXO1 in hdLECs stimulated with combined SVEP1 and ANG2 compared to ANG2 or SVEP1 alone (Figure 3C), indicating that ANG2 potentiates the activation of PI3K-Akt and thus inactivates FOXO1. ANG2 by itself had a comparable effect on the FOXO1 inactivation as SVEP1, with both proteins showing a significant effect (Figure 3F).

**Figure 3:**
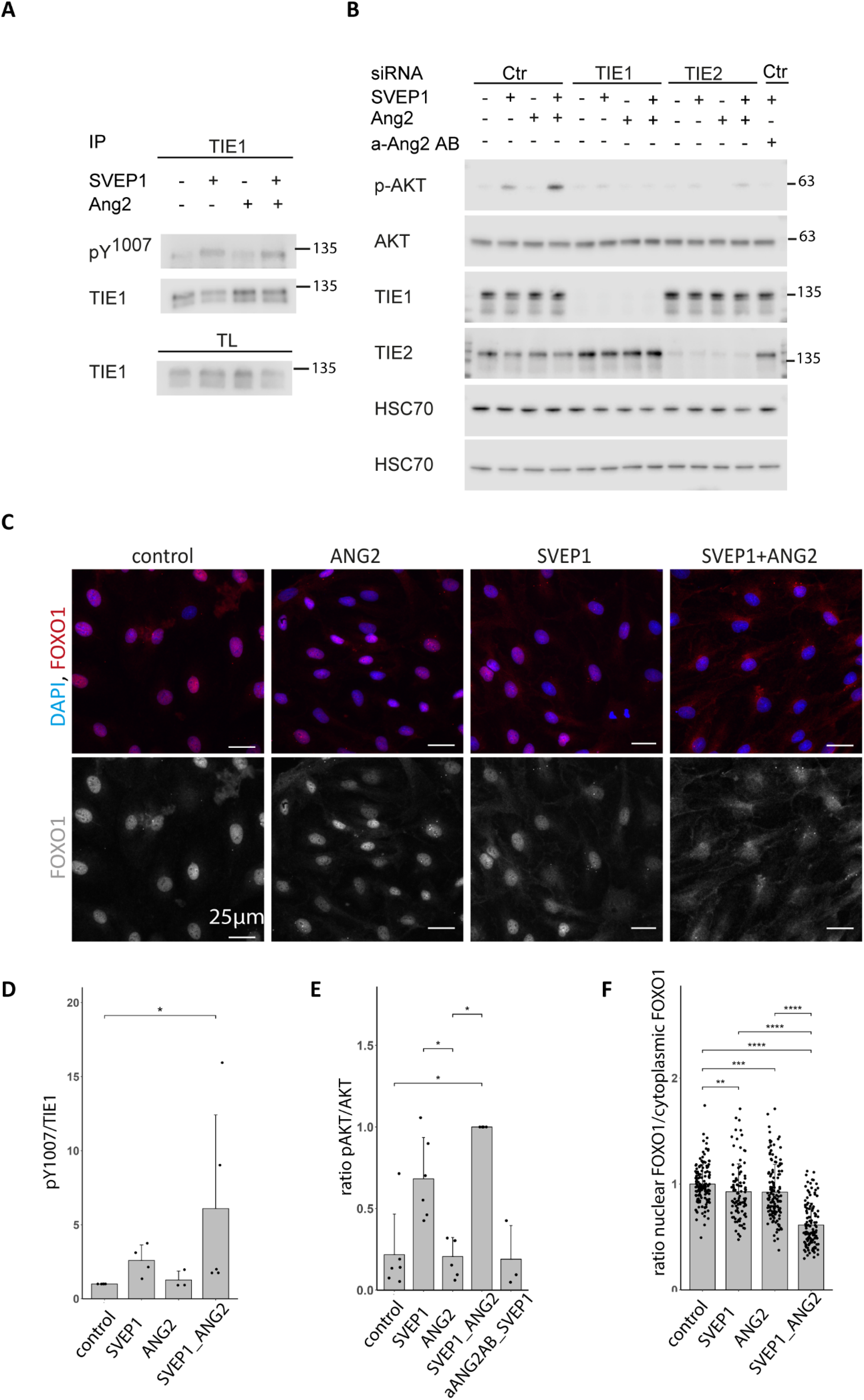
Simultaneous stimulation of LECs with SVEP1 and ANG2 leads to phosphorylation of TIE1 and AKT as well as nuclear exclusion of FOXO1 *in vitro*. (**A**) Western blot analysis of TIE1 phosphorylation at Y^1007^ residue in SVEP1 and/or ANG2 stimulated LECs after immunoprecipitation of TIE1. TIE1 is phosphorylated after simultaneous stimulation with SVEP1 and ANG2. The antibody anti–phospho-TIE2 (pY992, AF2720 R&D Systems) detects both, TIE1 phosphorylation (at Y1007 residue) and TIE2 phosphorylation (at Y992 residue), but can be distinguished by size (Brouillard et al. 2024). (**B**) Western blot analysis of p-AKT and AKT in the cell lysate of SVEP1 and ANG2 stimulated LECs. Knock-down of either TIE1 or TIE2 leads to reduced induction of phosphorylation of AKT by SVEP1-ANG2 (**C**) FOXO1 (red) and DAPI (blue) staining of LECs stimulated with only SVEP1 or ANG2, or in combination. FOXO1 staining is restricted to the nucleus of unstimulated (control) LECs, whereas in cells stimulated with SVEP1 and ANG2 together FOXO1 is in the cytoplasm. (**D**) Quantification of the pTIE1/total TIE1 ratios shown in A. (N = 5; *p adj = 0,045; Mann–Whitney U test; Values are presented as means ± SD) (**E**) Quantification of the pAKT/total AKT ratios shown in B. Values are presented as means ± SD, N = 3/6; Mann–Whitney U test; Control vs. ANG2_SVEP1 p.adj = 0.028 *; SVEP1 vs. ANG2 p.adj = 0.022 *; ANG2 vs. ANG2_SVEP1 p.adj = 0.028 *; ANG2_SVEP1 vs. aANG2AB_SVEP1 p.adj = 0.091. (**F**) The ratio of nuclear FOXO1/cytoplasmic FOXO1 decreases after stimulation with either ANG2 or SVEP1. Stimulation of LECs with simultaneous SVEP1 and ANG2 leads to a further reduction of the nuclear FOXO1/cytoplasmic FOXO1 ratio. Each datapoint represents the mean value per image of the ratio of nuclear to cytoplasmic FOXO1 per cell (Mann–Whitney U test; Values are presented as means ± SD; N=4).

### SVEP1, TIE1, and ANG1/2 are predicted to form higher-order complexes

The results show that SVEP1 together with ANG2 lead to phosphorylation of TIE1, which then activates the PI3K-AKT signaling pathway, leading to nuclear exclusion of FOXO1. Previous studies have demonstrated the importance of TIE2 receptor clustering and its impact on downstream signaling (Kim et al. 2009). To explore the ability of the SVEP1, TIE1, and ANG1/2 complexes to form higher order assemblies, we utilized AF3 to model a 2:2:2 complex. AF3 predicted that the previously modelled ternary complexes (Figure 2A) are incorporated into a larger assembly of dimerized TIE1 receptors (Figure 4A). In the presence of ANG1/2 and SVEP1, AF3 predicts that TIE1 dimerization is mediated by Fn3-Fn2’ contact as well as crossing over at the D3 and D4 domains (Figure 4B) with acceptable confidence metrics (Supplementary Figure 4 and Supplementary Figure 4.1). In addition to TIE1 dimerization, both ANG1 and ANG2 dimerize via a N-terminal coiled-coil domain and can multimerize into larger assemblies by interchain disulfide bonds (Kim et al. 2005). ANG1/2 multimerization is crucial for proper TIE2 activation (Kim et al. 2009) and ANG1/2 together with SVEP1 may have a similar effect with TIE1, clustering TIE1 receptors, even if TIE1 does not dimerize on its own (Figure 4C). Both proposed models would position TIE1 receptors in close spatial proximity, thereby increasing the cytosolic signaling strength of the TIE1 kinase domains upon binding of SVEP1 and ANG1/2. While the models put forth are of good quality, further experimentation is needed to confirm the structural model of SVEP1, TIE1, and ANG1/2 multimerization proposed.

**Figure 4:**
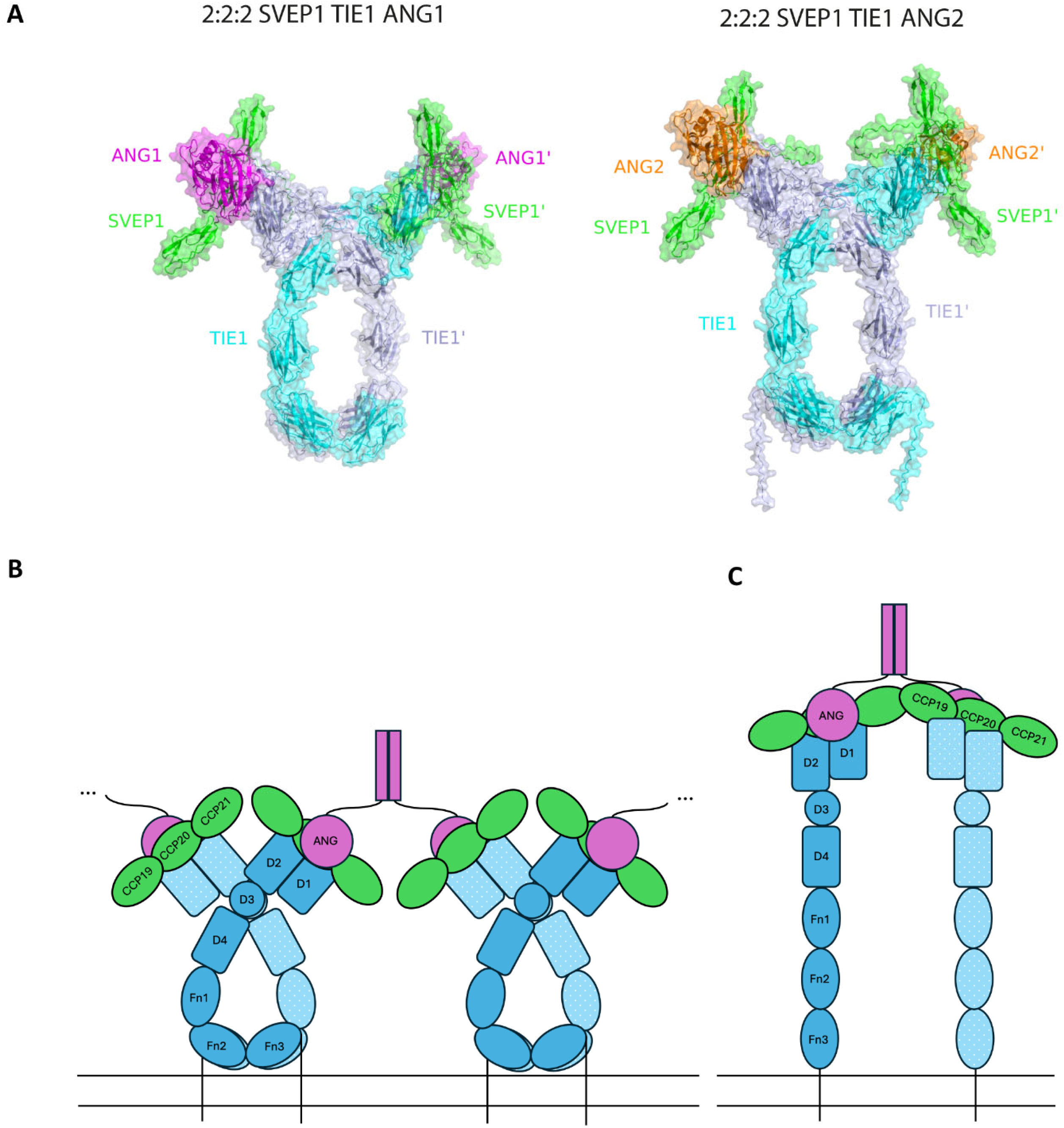
SVEP1, TIE1 and ANG1/2 can form 2:2:2 complexes mediated by SVEP1. (**A**) Larger hexameric complexes of SVEP1-TIE1 D1-D4-ANG1/2 in a 2:2:2 stoichiometry could be confidently modeled with AF3. They show a symmetric assembly linked by predicted TIE1 dimerization sites at FN3-FN2’ as well as the D3 and D4 domains. Similar to the 1:1:1 complex of SVEP1-TIE1-ANG1/2, ANG1/2 makes contact with the D1 and D2 domains of TIE1 by inclusion of the SVEP1 chain (CCP19-21). (**B**) Schematic model of SVEP1 enabling the multimerization of TIE1 receptors on the cell surface by cross-lacing of TIE1 dimers and N-terminal coiled-coil dimers (or greater assemblies) of ANG1/2. (**C**) Schematic model of TIE1 clustering via only ANG1/2 mediated multimerization.

## Discussion

TIE1 has long been considered an orphan receptor. Recently, we and others have shown that *svep1* and *tie1* genetically interact (Hußmann et al. 2023) and that SVEP1 binds to murine and human TIE1 (Hußmann et al. 2023; Sato-Nishiuchi et al. 2023). Here, we extend the analysis of SVEP1 as a binding ligand for TIE1 with the assistance of AF3 modeling and experimental support that demonstrates potentiated recruitment of TIE1 by SVEP1 in the presence of angiopoietins.

It is known that TIE2 is activated through binding of angiopoietins, including ANG1 and ANG2 (Davis et al. 1996; Maisonpierre et al. 1997), and that in humans and mice, angiopoietins do not directly bind to TIE1. In zebrafish, however, Ang1 binds to Tie1, Tie2 mutants do not have an obvious cardiovascular phenotype, and the *tie2* locus has been lost in many teleost species (Jiang et al. 2020). Therefore, zebrafish differ from humans and mice in terms of the TIE pathway, as zebrafish appear to exclusively use Tie1 and not Tie2 (Gjini et al. 2011; Jiang et al. 2020; Morooka et al. 2024). Both SPR testing and AF3 modeling revealed that SVEP1 (CCP15-CCP24) does not interact with TIE2. This is in line with trans-well migration assays, which revealed that SVEP1 induces the migration of lymphatic endothelial cells through binding to TIE1, but not TIE2 (Sato-Nishiuchi et al. 2023). Whether TIE2 is able to bind SVEP1 in a condition where other proteins like angiopoietins are present is still a question of interest and requires further investigation.

We here demonstrate via AF3 model and co-immunoprecipitation, that the affinity of TIE1 for SVEP1 is highly enhanced by the simultaneous binding of the C-terminal domains of ANG2 to the CCP20 module, thus generating a complex of SVEP1, angiopoietins, and TIE1 molecules in which the angiopoietins are also able to bind TIE1. Utilizing AF3 modeling, we were able to confidently predict an interaction hub with a 2:2:2 complex of SVEP1, TIE1, and ANG1/2, where ANG1/2 binds different SVEP1 molecules by cross-lacing of N-terminal coiled-coil dimers, which then brings several TIE1 receptors into spatial proximity on the cell surface. This could lead to a cumulative increase in the signal strength of the cytosolic kinase TIE1. While TIE2 has been shown to dimerize via Fn3-Fn3’ contacts (Moore et al. 2017), evidence of TIE1 dimerization is limited. Leppanen et al. (2017) captured a strand swapped dimer of TIE1 Fn3 by X-ray crystallography (pdb: 5N06). However, solution SAXS data indicated monomeric TIE1 Fn3, leading to the conclusion that the observed crystallography dimer is likely to be a crystal artifact. However, our model as well as experimental data predicts that in presence of SVEP1 and ANG1, TIE1 can dimerize and thus is phosphorylated upon ligand binding. So far, phosphorylation of TIE1 has not been detected when cells were stimulated with either SVEP1 or ANG2 only (Saharinen et al. 2005; Sato-Nishiuchi et al. 2023; Savant et al. 2015). Sato-Nishiuchi et al. (2023) could not detect an increase in the baseline phosphorylation of murine TIE1 in either LECs or 293-F cells that were treated with Svep1. Only TIE2 co-expression increased TIE1 phosphorylation irrespective of the presence or absence of Svep1. In blood and lymphatic endothelial cells, only COMP-Ang1 and ANG1 have been shown to stimulate the phosphorylation of TIE1 (Saharinen et al. 2005; Savant et al. 2015; Yuan et al. 2007). Co-transfection of TIE1 and TIE2 enhances TIE1 activation and phosphorylation due to heteromeric TIE1-TIE2 complexes. On the contrary, TIE2 phosphorylation is not enhanced by the presence of TIE1 (Saharinen et al. 2005), and TIE1 is not phosphorylated in the absence of TIE2 upon ANG1 stimulation (Savant et al. 2015). According to current knowledge, the phosphorylation of TIE1 is therefore dependent on TIE2. In zebrafish, however, it was shown that Tie1 can be phosphorylated (Morooka et al. 2024) and that Tie1, not Tie2, is the relevant receptor for lymphangiogenesis during early development. Furthermore, it was reported that autophosphorylation of TIE1 (Y1007) is not dependent on TIE2 but on TIE1, because substitutions in the kinase domain of TIE1 result in a loss of its baseline phosphorylation (Brouillard et al. 2024). Additionally, after COMP-Ang1 stimulation of TIE1-transfected 293T cells that do not express TIE2, weak phosphorylation is observed, whereby the activation is mediated by ANG1 and not by the COMP domain, a minimal coiled-coil segment that oligomerizes the ANG1 C-term module (Saharinen et al. 2005), which suggests that the phosphorylation is not dependent on TIE1/TIE2 heterodimerization. Nevertheless, since TIE1 was considered an orphan receptor during that time, the result could not be explained in 2005, and it was speculated whether COMP-Ang1–induced Tie1 activation could involve complex formation with additional molecules.

Our results show phosphorylation of TIE1 after combined SVEP1-ANG2 stimulation of hdLECs. SVEP1 is not expressed by HUVECs and LECs, but in non-endothelial cells (such as fibroblasts) in close proximity to LECs (Karpanen et al. 2017; Hußmann et al. 2023; Wang et al. 2020). According to data from the human protein atlas (https://www.proteinatlas.org), SVEP1 is also not expressed in HEK293 cells. Since SVEP1, which is necessary for the phosphorylation of TIE1, is provided by cells in the immediate vicinity of ECs and not by ECs themselves, the phosphorylation of TIE1 after ANG2 stimulation was never detected in cell culture experiments without SVEP1 being present. Whether TIE2 is necessary for the phosphorylation of TIE1 after SVEP1-ANG2 treatment needs further investigation. Our model shows that after binding of SVEP1 to TIE1, ANG1/2 are then able to dock onto this complex, and serve as a ‘molecular glue’ for both SVEP1 and TIE1. Thus, TIE1-bound SVEP1 would serve as a cell-surface-tethered interaction hub for angiopoietins that are oligomerized – by either the native coiled-coil extensions or the minimal COMP domain – and create cross-linked fields of signaling TIE1 receptors (Figure 4).

The PI3K/Akt signaling pathway is involved in Svep1-induced LEC migration, as LECs show an increased Akt phosphorylation after Svep1 treatment (Sato-Nishiuchi et al. 2023). We were able to show that combined SVEP1-ANG2 stimulation leads to an even higher phosphorylation of AKT in human LECs than the sum of SVEP1 or ANG2 stimulations alone. Blocking of ANG2 using a function blocking antibody prevents AKT phosphorylation after stimulation with SVEP1, implying that SVEP1 cannot induce downstream signaling on its own, but only in combination with angiopoietins.

Furthermore, both TIE1 and TIE2 are required for the full signaling potential of SVEP1-ANG2, as knock-down of either TIE1 or TIE2 reduced the phosphorylation of AKT upon SVEP1-ANG2 stimulation. A signaling event downstream of PI3K/Akt is the inactivation and nuclear exclusion of FOXO1 (Daly et al. 2004), which is reduced in Tie1-silenced vascular endothelial cells (Korhonen et al. 2016). Furthermore, SVEP1 induces FOXO1 nuclear exclusion. This was shown in LECs as well as *in situ* where FOXO1 immunostaining of lymphatic vessels in wild type mice was detected in the nucleus, whereas in *Svep1*-deficient mice, FOXO1 staining was detectable in both the nucleus and cytoplasm (Sato-Nishiuchi et al. 2023). During secondary sprouting of endothelial cells in the zebrafish trunk, Tie1 induces nuclear exclusion of *foxo1a,* since *tie1* mutants displayed an enhanced localization of *foxo1a* in the nucleus (Morooka et al. 2024). Thus, the nuclear versus cytoplasmic ratio of FOXO1 is used as a readout for signaling in a number of systems. Here we show that stimulation of LECs with a combination of SVEP1 and ANG2 leads to a stronger nuclear exclusion of FOXO1 than with either factor alone, thus indicating that stronger binding of SVEP1 to TIE1 also potentiates downstream signaling of TIE1.

The CCP20 module of SVEP1 acts as a critical binding partner for TIE1, but not TIE2, when interacting with ANG1/2. However, it remains unclear whether TIE2 could bind to other sites of SVEP1 in the presence of other proteins, since in the SPR assay and the immunoprecipitation assays, only part of the C-terminus of the SVEP1 protein was used. Also, TIE1 but not TIE2 is required for SVEP1-induced migration (Sato-Nishiuchi et al. 2023), but both are necessary for SVEP1-stimulated phosphorylation of AKT. Furthermore, in capillaries, only depletion of *Tie1*, but not of *Tie2,* had an effect, whereas in collecting vessels, depletion of *Tie2* resulted in a phenotype (Korhonen et al. 2022). This result can most likely be explained by high expression levels of TIE2 in collecting vessels, but low expression levels of TIE2 in capillaries. Additionally, it is proposed that SVEP1 acts as a modifier of TIE2 expression during Schlemm’s canal development as patients that are trans-heterozygous for TIE2 and SVEP1 develop glaucoma and the p.R997C-SVEP1 variant, which is mutated in some glaucoma patients and located shortly after the Furin cleavage site, fails to enhance TIE2 expression in HEK293T cells compared to wild type SVEP1 (Young et al. 2020). Depending on the downstream signaling event as well as the lymphatic cell type (capillaries versus collecting vessels), TIE2 might be needed for phosphorylation of TIE1 either independently of SVEP1 or for binding to other domains of SVEP1 than CCP19-21.

The activation mechanisms of the TIE receptors have long been a point of contention. Especially TIE1 activation and how possible signaling occurs has remained enigmatic, and only recently SVEP1 has been demonstrated to serve as a ligand for the TIE1 transmembrane receptor, which for many years had been considered an orphan receptor with phosphorylation believed to be dependent on TIE2. We here show that SVEP1-ANG2 is able to cause phosphorylation of TIE1 and show that the presence of ANG2 results in stronger recruitment of TIE1 to SVEP1, supporting a model that predicts a higher order complex of SVEP1, TIE1, and ANG1/2. In such complexes, ANG1/2 binds different SVEP1 molecules by cross-lacing of N-terminal coiled-coil dimers, which then brings several TIE1 receptors into spatial proximity on the cell surface. This could lead to formation of a signaling hub, where the signal strength of the cytosolic kinase TIE1, or another signaling moiety, is increased.

## Material and Methods

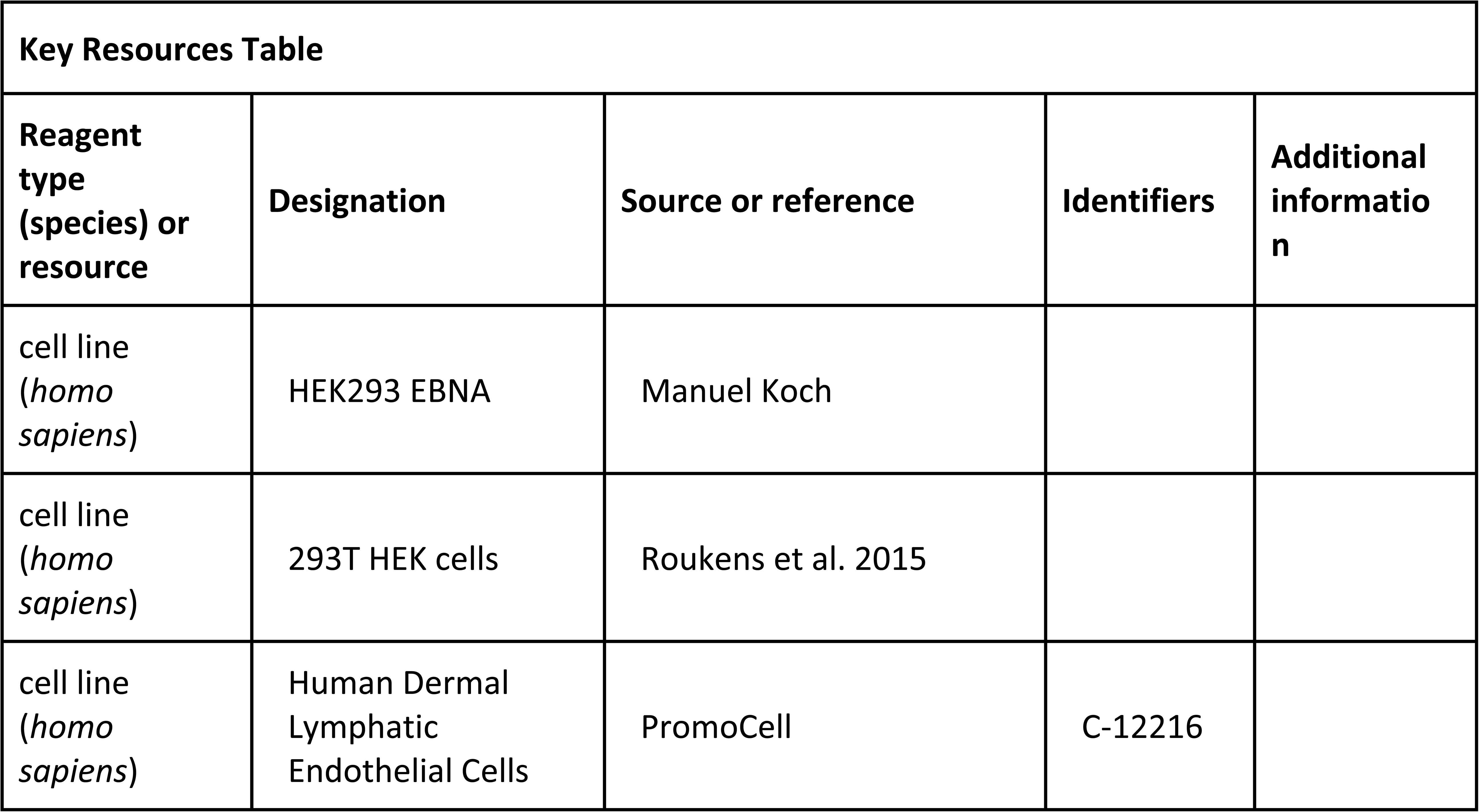
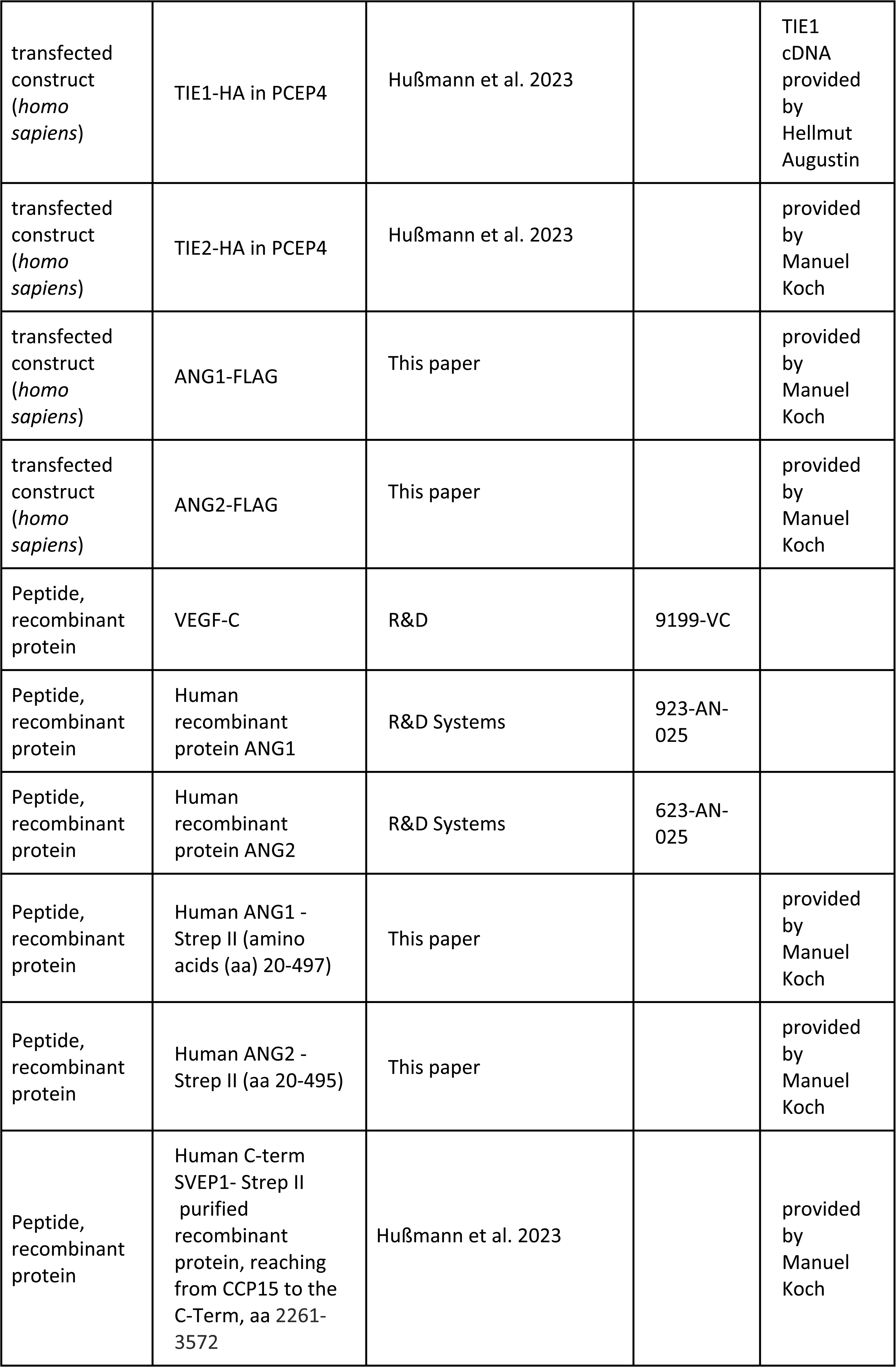
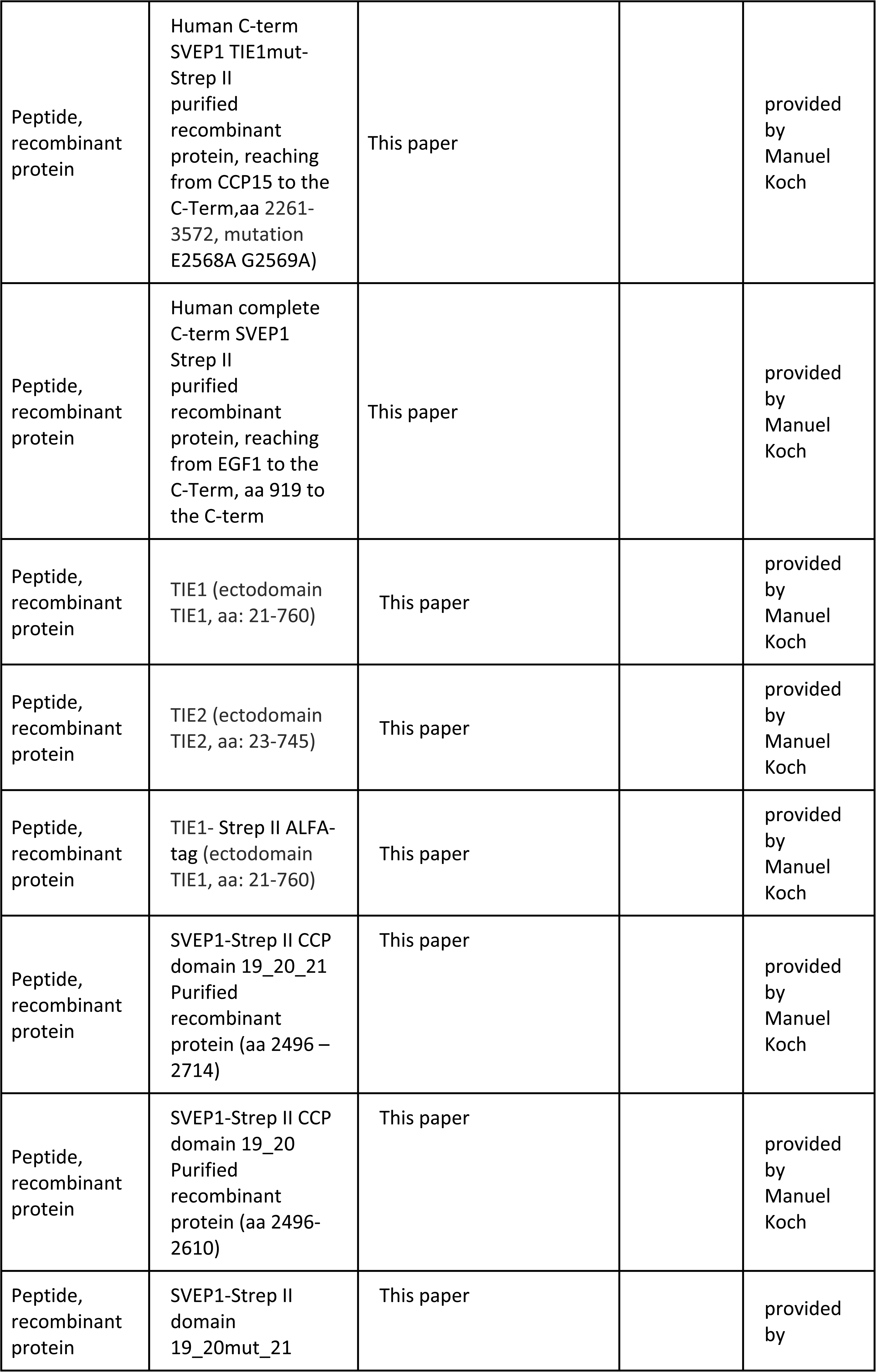
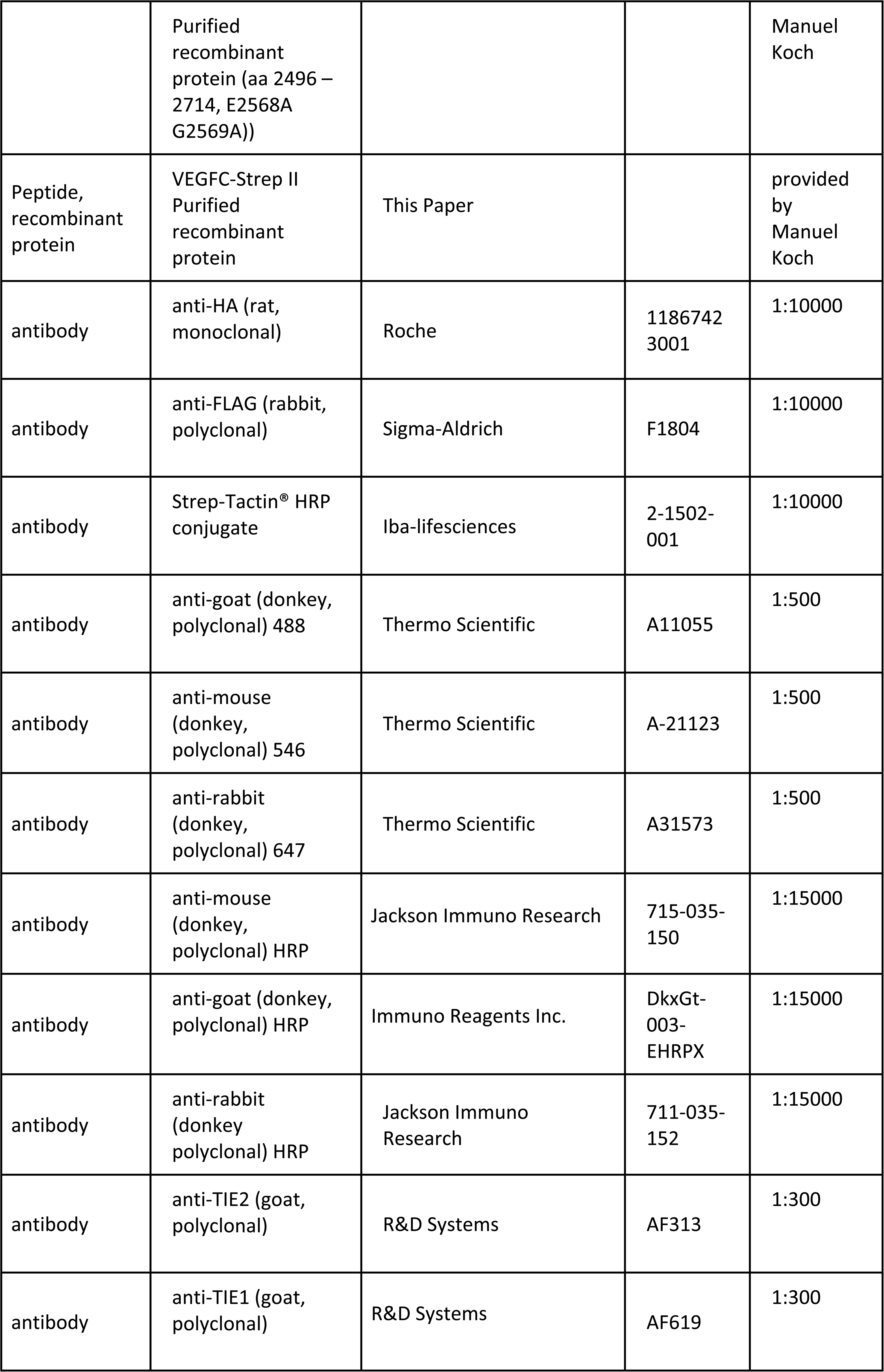
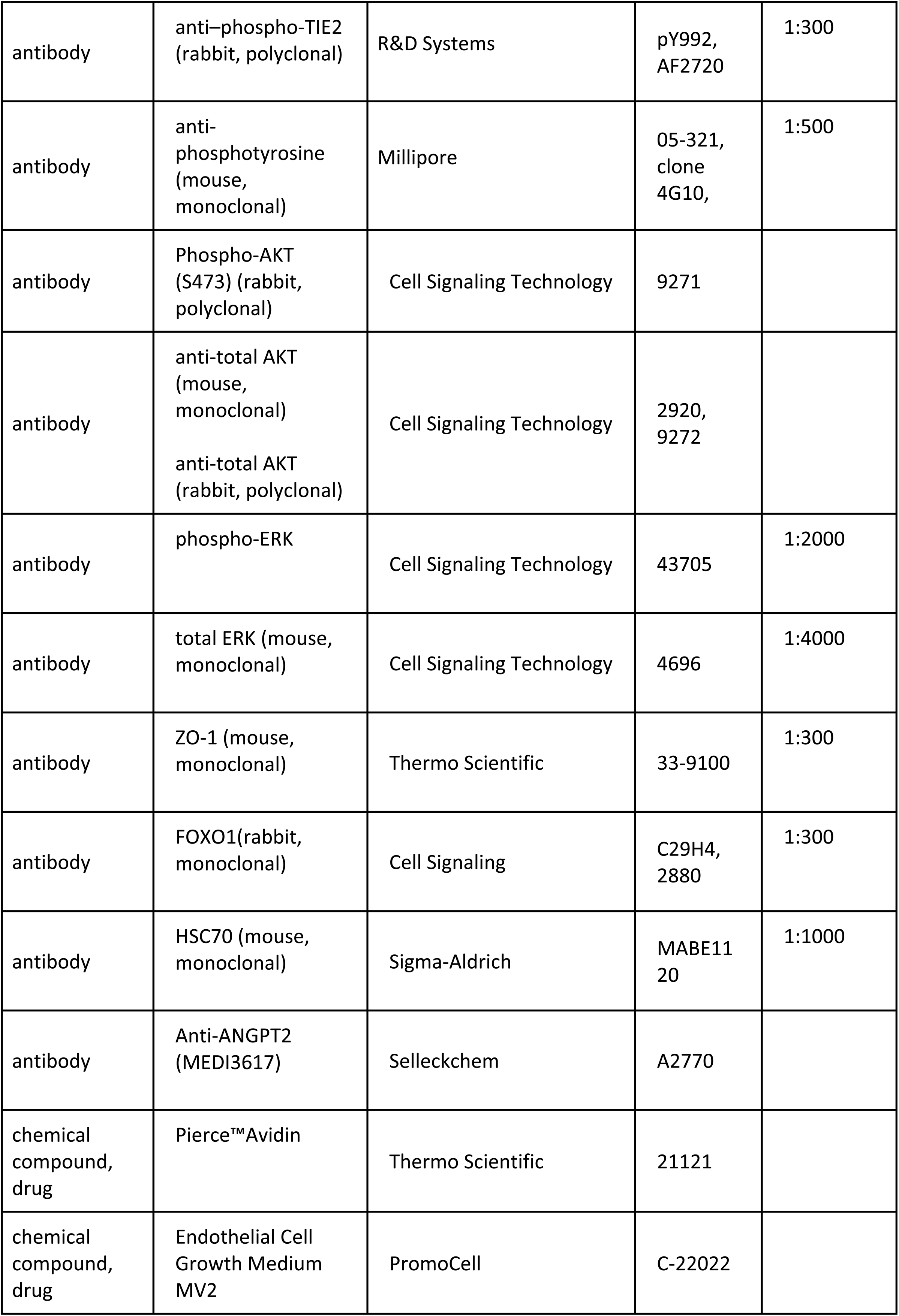
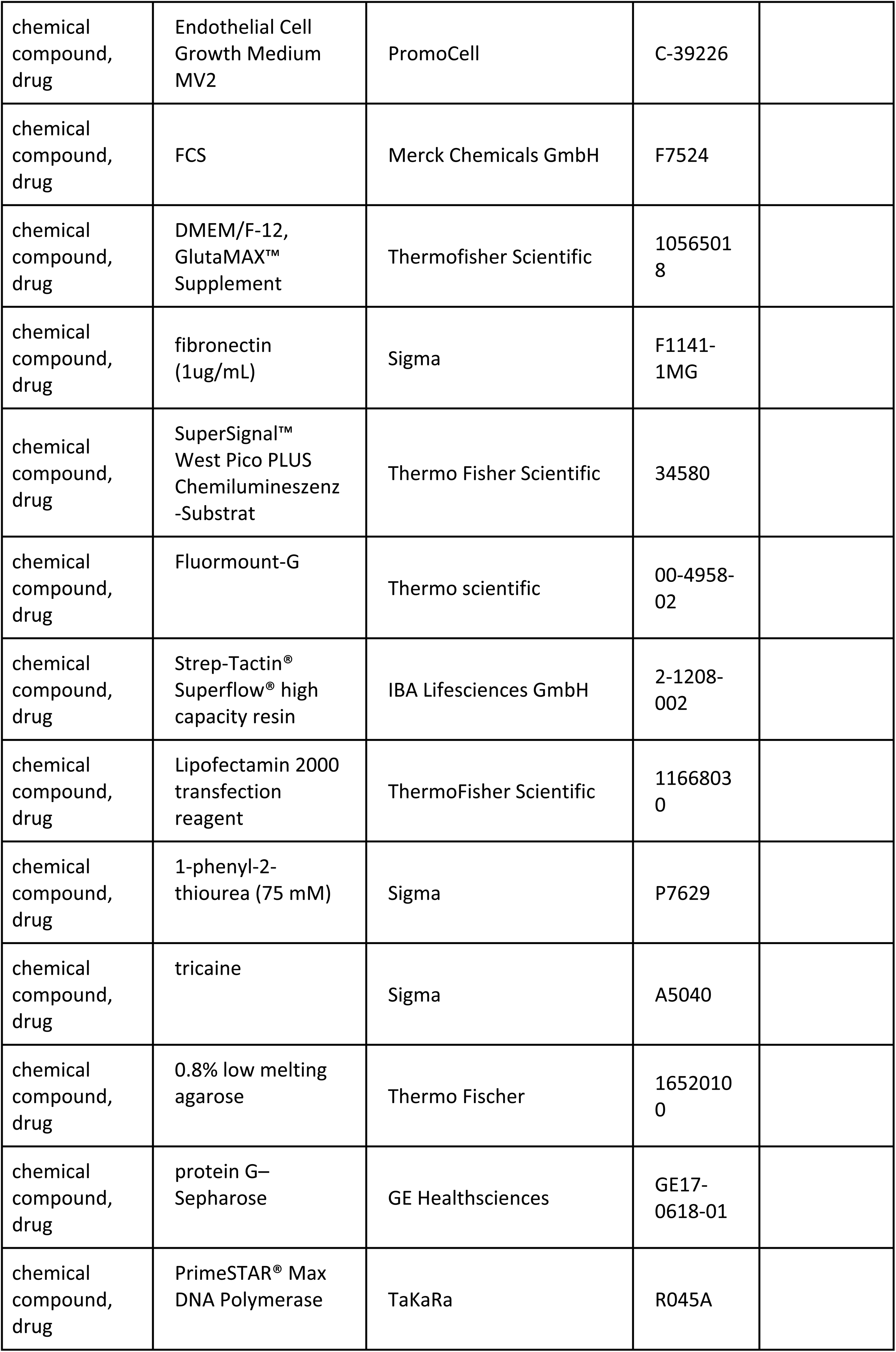
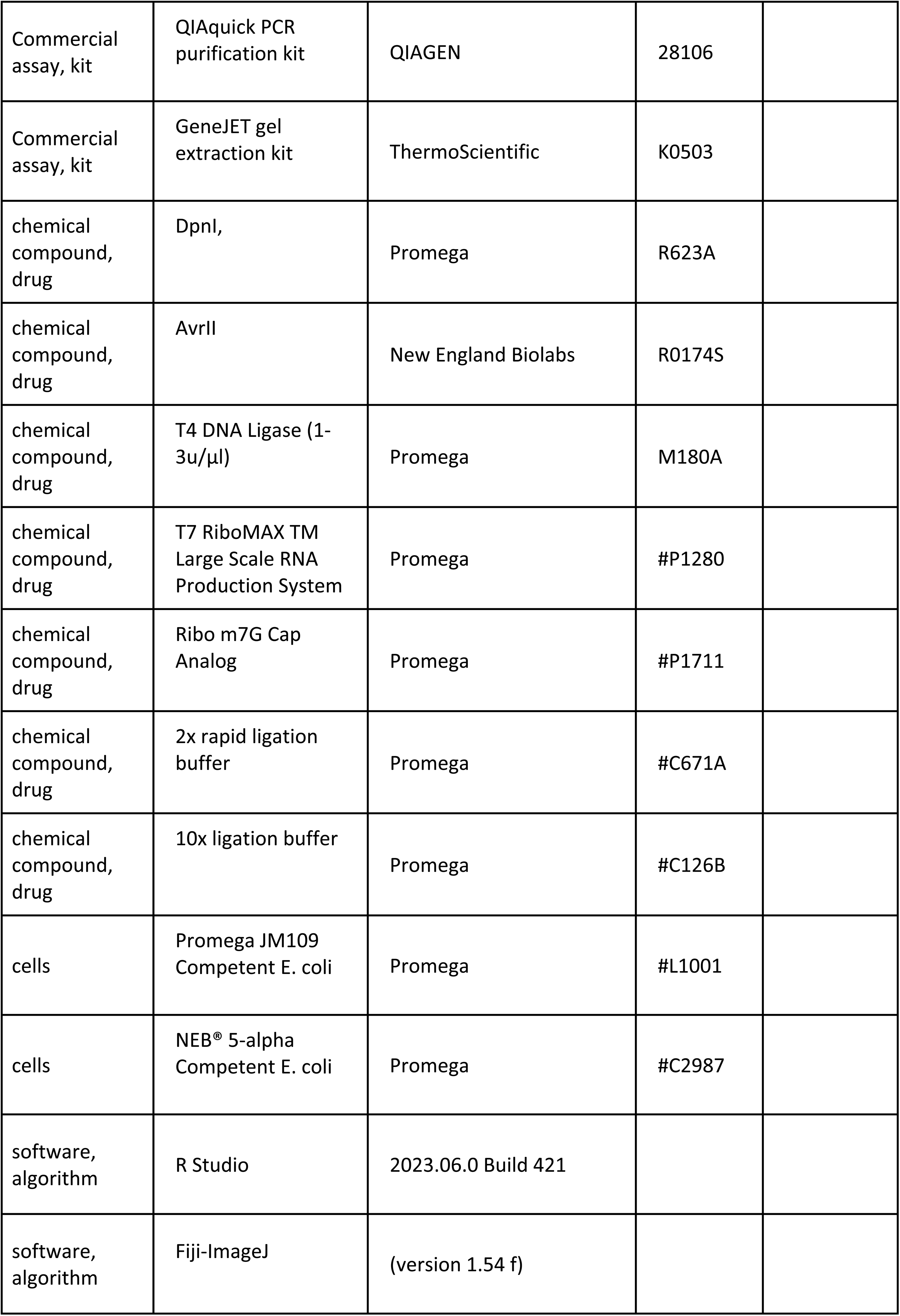
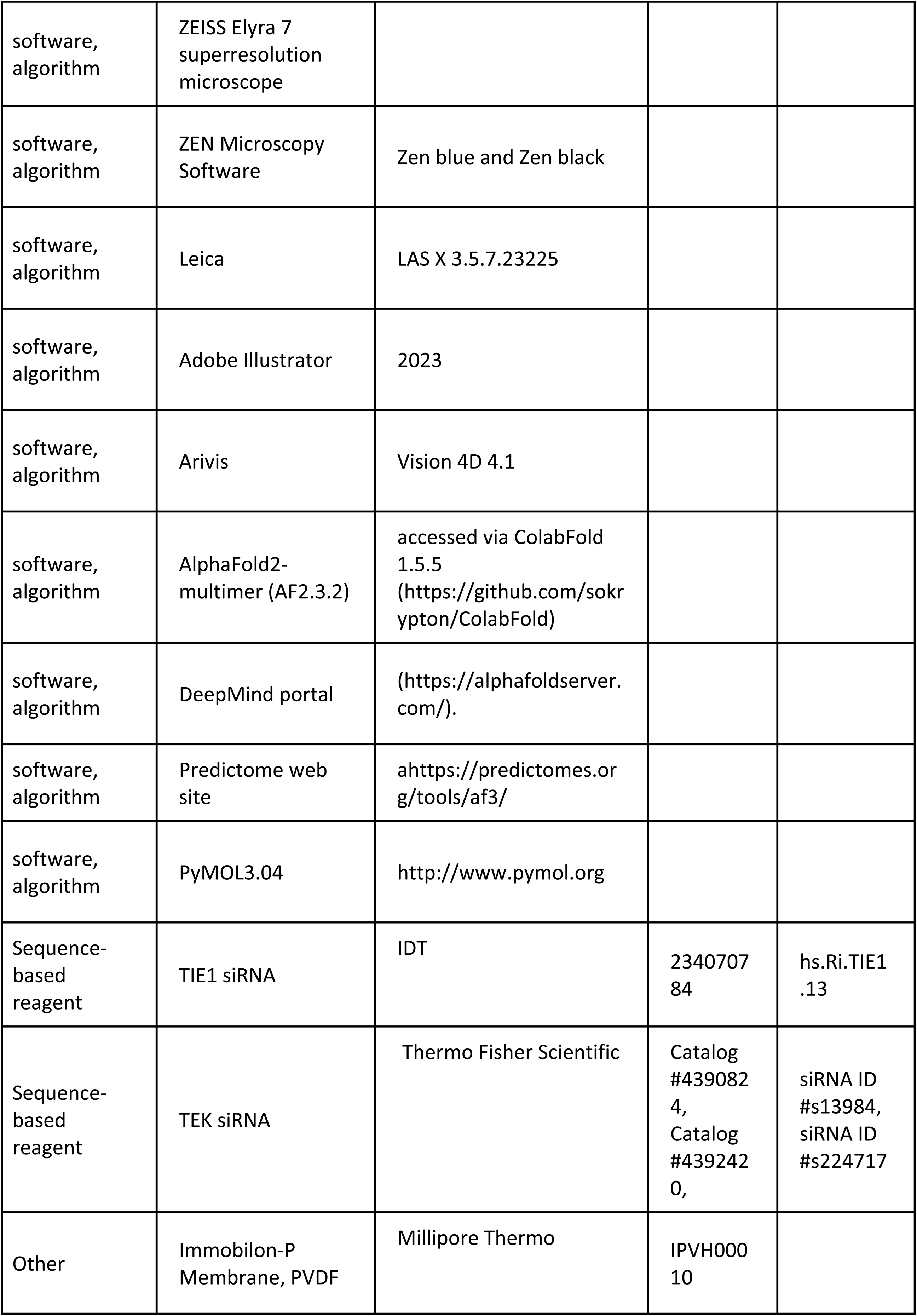

### Recombinant protein expression

Recombinant proteins (SVEP1, GenBank: NM_153366.5), (ANG1, GenBank: NM_001199859), (ANG2, GenBank: NM_001118887), TIE1 (GenBank: NM_005424) and TIE2 (GenBank: NM_000459) were amplified from human cDNA by PCR. Amino acids contained in the different recombinant proteins are listed in the key resource table. Protein production was carried out essentially as described (Hußmann et al. 2023; Meier et al. 2023). Briefly, SVEP1, TIE1 and ANG2 containing an N-terminal twin Strep-tag were expressed in HEK 293 T cells using the inducible sleeping beauty transposon system and secreted into the medium. Tagged proteins were bound to Strep-Tactin XT 4Flow matrix (IBA Lifescience) equilibrated in 50 mM Tris pH 8.0, 150 mM NaCl. After washing with 50 mM Tris pH 8.0, 1 M NaCl proteins were recovered using Biotin containing BXT elution buffer (IBA Lifescience) and subsequently dialyzed into PBS. PCR products were cloned into a modified sleeping beauty transposon expression vector containing a BM40 signal peptide sequence and a Twin-Strep-tag. For recombinant protein production, stable HEK293 EBNA cell lines were generated employing the sleeping beauty transposon system (Kowarz et al. 2015). Briefly, expression constructs were co-transfected with a transposase plasmid (1:10) into the HEK293 EBNA cells using FuGENE HD transfection reagent (Promega). After selection with puromycin (2 µg/ml), cells were induced with doxycycline. Supernatants were filtered and the recombinant proteins purified via Strep-Tactin®XT (IBA Lifescience) resin. Proteins were then eluted by biotin-containing TBS-buffer (IBA Lifescience) and subsequently dialyzed into PBS. The human sequences of *ANGPT1* (corresponding to NM_001199859, aa: 20 – 497) and *ANGPT2* (corresponding to NM_001118887, aa: 20–495) were cloned into the PCEP episomal expression system (transient) including an N-terminal Flag-tag sequence.

### Surface Plasmon Resonance

Protein-protein interactions were analyzed by surface plasmon resonance (SPR) experiments using a Biacore T200 system (Cytiva, Uppsala, Sweden). Human active SVEP1-Strep II recombinant protein, reaching from CCP15 to the CCP24, aa 2261-2890 and TIE1 (ectodomain TIE1, aa: 21-760) or TIE2 (ectodomain TIE2, (aa: 23-745) were used. For immobilization of hSVEP1, a CM5 sensor chip was activated with N-hydroxysuccinimide (NHS) and 1-ethyl-3-(3-dimethylaminopropyl) carbodiimide hydrochloride (EDC) according to the manufacturer’s instructions, and 2050 response units (RU) of SVEP1 were coupled in 10 mM sodium acetate buffer, pH 4.0, at a flow rate of 10 µL/min. For the interaction measurements, the analytes, TIE1 and TIE2, were passed over the sensor chip at a flow rate of 30 µl/min in running buffer (10 mM HEPES, 150 mM NaCl and 0.005% Tween20). The concentrations for the TIE proteins were 200 nM, 100 nM, 50 nM, 25 nM, 12.5 nM, 6.25 nM and 3.125 nM. The binding kinetics were calculated by fitting a 1:1 interaction model using the BIAevalution software (Cytiva, Uppsala, Sweden).

### Negative staining

The interaction of the SVEP1 C-terminus with Angiopoietin 1 was visualized by negative staining and transmission electron microscopy. Briefly, the proteins were incubated for 1 hour at 37°C in Tris-buffered saline (TBS), pH 7.4. Prior to the visualization in the electron microscope, the proteins were negatively stained with 1% uranyl acetate on the grids. In some experiments, the proteins were directly labelled with 5 nm (Sigma-Aldrich 752568) and 10 nm (Sigma-Aldrich 752584) colloidal gold, respectively, as previously described (Baschong and Wrigley 1990). Specimens were examined in a Philips/FEI CM 100 TWIN transmission electron microscope operated at 60 kV accelerating voltage. Images were recorded with a side-mounted Olympus Veleta camera with a resolution of 2048 × 2048 pixels (2k × 2K) and the ITEM acquisitions software.

### ELISA binding assays

ELISA assays were performed as described (Eble 2018; Meier et al. 2023). Briefly, SVEP1 or SVEP1 mutants (2 µg/ml in TBS) were coated on 96 well plates (Nunc MaxiSorp, 50 µl/well) for 1 h at room temperature and subsequently washed with TBS. Plates were blocked using Pierce Protein-Free Blocking Buffer (ThermoFisher Scientific) for 1 h at room temperature followed by 1 % BSA in TBS over night at 4°C. Bound SVEP1 was overlaid with serial dilutions of TIE1 in the presence or absence of ANG2, or TIE mutant containing an ALFA tag, or biotinylated ANG2 protein, and incubated for 1 h at room temperature. After washing with PBS, protein complexes were fixed with 2.5 % glutaraldehyde in PBS for 10 min and the reaction stopped by washing with TBS. Bound TIE or ANG2 proteins were detected with HRP conjugated anti ALFA-tag nanobody or anti-Biotin antibody and stained using 1-Step Ultra TMB-ELISA (ThermoFisher Scientific). After addition of 10% sulfuric acid, absorbance reading at 450 nm was done on a Tecan plate reader equipped with Magellan Pro software. Non-linear regression analysis of data was performed using GraphPad Prism 11 software.

### Cell culture and treatment/stimulation

HEK293T cells were cultured in DMEM (Thermo Fisher Scientific #61965026) and seeded one day before transfection utilizing Lipofectamine™ 2000 (Thermo Fisher Scientific, #11668027). hdLECs (Human Dermal Lymphatic Endothelial Cells, PromoCell) were cultured in Endothelial Cell Growth Medium MV2 (PromoCell, #C-22022) on Fibronectin-coated (Sigma, #F1141-1MG, 1ug/mL) culture flasks. LEC culture medium was supplemented with 25 ng/ml VEGF-C (R&D, #9199-VC). Cells were seeded on ibidi slides (ibidi, #81201) and grown to confluency before the treatment.

For knock-down experiments, hdLECs were transfected with *TIE1*, *TIE2* or control siRNA (see key resources table) using Lipofectamine™ RNAiMAX Transfection Reagent one day after splitting and two days prior to analysis of AKT phosphorylation.

For the pull-down assay, different versions of human StrepII tagged SVEP1 protein and a StrepII tagged control protein were incubated with 25 µl Strep-TactinXT 4Flow high-capacity resin (iba-lifesciences, #2-5030-002) in 500 µl binding buffer (50 mM Tris–HCl at pH 7.5, 100 mM NaCl, 0.02% Triton X-100) for 30 min. Subsequently, the cell lysate of TIE1-HA transfected HEK293T cells was added. After 2 hours of incubation, the beads were washed 5 times with Ripa buffer (50 mM Tris (pH 7.5), 1% NP-40, 0,1% SDS, 0,5% Na-deoxycholate, 150 mM NaCl) and boiled for 5 min at 95°C in sample buffer. For pull down assays that include angiopoietins, TIE1-HA and ANG2-FLAG/ANG1-FLAG were co-transfected and the cell lysates were pre-cleared with Strep-TactinXT 4Flow high-capacity resin, before adding them to the StrepII tagged SVEP1 protein.

For TIE1 autophosphorylation, hdLECs were starved for 3 hours in 0.5% BSA-containing MV2 medium, followed by 1 hour of stimulation with ANG2 and SVEP1 alone or in combination. The cells were lysed in PLCLB lysis buffer (150 mM NaCl, 5% glycerol, 1% Triton X-100, 1.5 mM MgCl2, 2mM CaCl2, 50 mM HEPES, pH 7.5, 1 x Complete-EDTA-free proteinase inhibitors) followed by TIE1 immunoprecipitation using anti-TIE1 antibodies (AF619, R&D Systems) and protein G–Sepharose (GE Healthsciences Ab, #GE17-0618-01) at +4°C. After 2 hours of incubation, the beads were washed 5 times with PLCLB lysis buffer and boiled for 5 min at 95°C in sample buffer.

For analysis of pAKT, growth medium of confluent hdLECs was changed to MV2 medium containing growth supplements but lacking VEGFC. Cells were starved for 3 hours in 0.5% BSA-containing MV2 medium, followed by 40-60 minutes of stimulation with ANG2 (0.5 µg/mL) and/or SVEP1 (1 µg/mL). Cells were treated with anti-ANG2 antibody (2 µg/ml; MEDI3617 A2770, Selleckchem) for 2 h prior to stimulation with SVEP1. The cells were lysed in lysis buffer (20 mM Tris-HCl (pH 7.4), 150 mM NaCl, 2mM CaCl2, 1.5 mM MgCl2, 1 mM Na3VO4, 1 % (v/v) Triton-X-100, 0.04 % (w/v) NaN3, 1 x Complete-EDTA-free proteinase inhibitors) or PLCLB lysis buffer and centrifuged before gel analysis.

### Western blot

Cell lysates and cell precipitates were subjected to Western blot analysis by separation in SDS-page and blotting onto PVDF membrane. Blots were probed with primary antibodies followed by HRP coupled secondary antibodies and ECL detection (SuperSignal™ West Pico PLUS Chemilumineszenz-Substrat, Thermo Fisher Scientific, 34580). Blots were imaged using an Odyssey FC (LICOR). Primary and secondary antibodies used for analysis are listed in the key resources table and are stated in the figure of the respective blot. Before using the Strep-Tactin® HRP antibody, the membrane was incubated in 2,5µg/ml Avidin (biotin blocking solution) for 10 minutes. HSC70 was used to assure equal loading of the gels and values were normalized by the level of HSC70.

### Immunocytochemistry and imaging

Before the treatment, hdLECs were starved for 4 hours in MV2 medium without supplements and with 0,5 % FBS. After treatment with SVEP1 and/or ANG2 (623-AN R&D Systems) for 1-hour cells were fixed with 4 % PFA for 10 minutes. Afterwards the fixed cells were permeabilized using 0,5% TritonX-100 in PBS for 5 minutes and blocked with 2% BSA for 1 hour. Incubation of the slides with the antibody was conducted over night at 4°C (ZO-1 (Thermo Scientific, 33-9100), FOXO1 (Cell Signaling, C29H4, #2880)). After washing the slides several times with 0,1%PBS-TritonX100 the secondary antibody staining was conducted at room temperature for 50 minutes (Donkey-anti-rabbit-647 (Thermo Scientific A31573), Donkey-anti-mouse-546 (Thermo Scientific), DAPI). Slides were washed and mounted using Fluormount-G (Thermo scientific 00-4958-02). Images were taken using ZEISS Elyra 7 super-resolution microscope in Apotome mode with 20x objective and processed with ZEN Microscopy Software (Zen blue 3.5 and Zen black). ZEISS arivis Software was used to calculate the pixel intensities inside the nucleus and in the cytoplasm. ZO1 was used to identify single cells and the ratio was calculated for each cell.

### Statistics and reproducibility

Data sets were tested for normality (Shapiro–Wilk) and equal variance. P-values of data sets with normal distribution were determined by Student’s *t*-test. In case data values did not show normal distribution, a Mann–Whitney U test was performed instead. For multiple comparison the p values were adjusted with Bonferroni correction. Results are presented as mean ± SD. Statistical tests were performed using GraphPad Prism 8 and R. For visualizations, the R package ggplot2 was used. All experiments were carried out at least three times. P values > 0.05 were considered not significant. * p≤0.05, ** p≤0.01, ***p≤0.001, **** p≤0.0001.

### Structure prediction and modeling of SVEP1 complexes

Amino acid sequences for human SVEP1 (https://www.uniprot.org/uniprotkb/Q4LDE5), TIE1/2 (https://www.uniprot.org/uniprotkb/P35590 and /Q02763) and ANG1/2 (https://www.uniprot.org/uniprotkb/Q15389 and /O15123) were retrieved from UniProtKB. The most recent version of AlphaFold2-multimer (v2.3.2) was accessed via ColabFold (v1.6.1) (https://github.com/sokrypton/ColabFold) while the DeepMind portal was used for AF3 (https://alphafoldserver.com/) modelling of >2500 amino acid jobs. For the template-free AF2.3 predictions, num recycles=auto and num models=5 were used, models were relaxed with AMBER, and ranked based on ipTM scores; in turn, default parameters were used for AF3. Top ranked model interfaces were analyzed and visualized with PDBePISA (Krissinel and Henrick 2007) and PyMOL v3.1.8 (The PyMOL Molecular Graphics System, Version 3.1.4.1, Schrödinger, LLC).

## Data availability

Scripts used for data analysis available at GitHub at https://github.com/MuensterImagingNetwork/Uphoff_et_al_2025/tree/main (copy archived at Münster Imaging Network, 2025). The datasets generated during and/or analyzed during the current study are available from the corresponding author on reasonable request.

## Acknowledgements

This work was supported by the CiM-IMPRS graduate school. We thank Sarah Weischer, Thomas Zobel and Jens Wendt (Münster Imaging Network, Cells in Motion Interfaculty Centre, University of Münster, Germany) for the support in imaging analysis. The Zeiss Elyra 7 microscope was funded by INST 211/901-1 FUGB.

---

**Supplementary figure 1.1:**
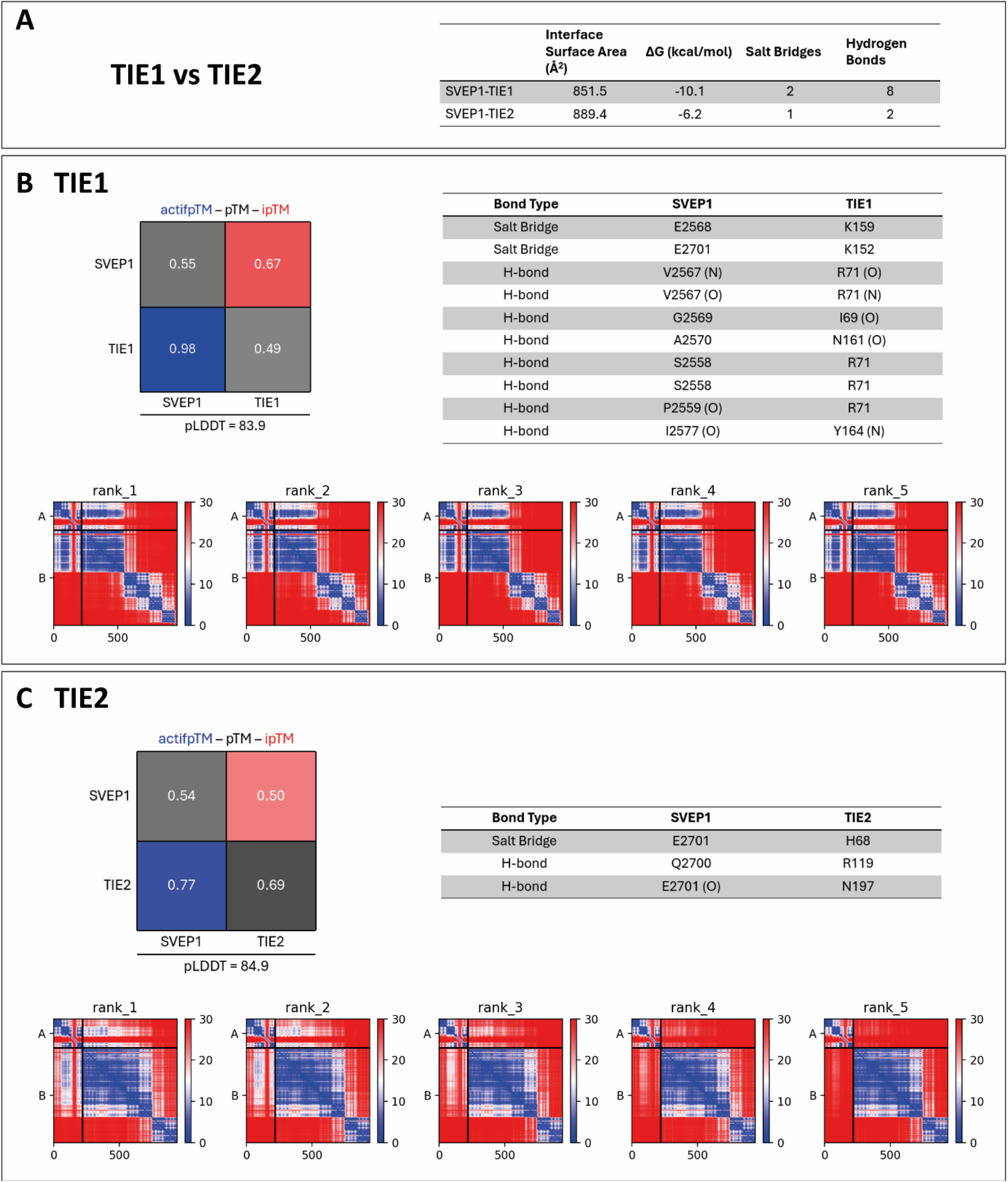
Statistics and (**A**) PDBePISA analysis of a SVEP1-TIE1ectodomain model vs a SVEP1-TIE2 ectodomain model depicting interface surface area, Gibbs free energy, salt bridges and hydrogen bond contacts. (**B**) and (**C**) statistics table, bonds and PAE plot of TIE1 (**B**) or TIE2 (**C**). PAE plot: A =SVEP1 CCP19-21, B=TIE1 or TIE2.

**Supplementary Figure 1.2:**
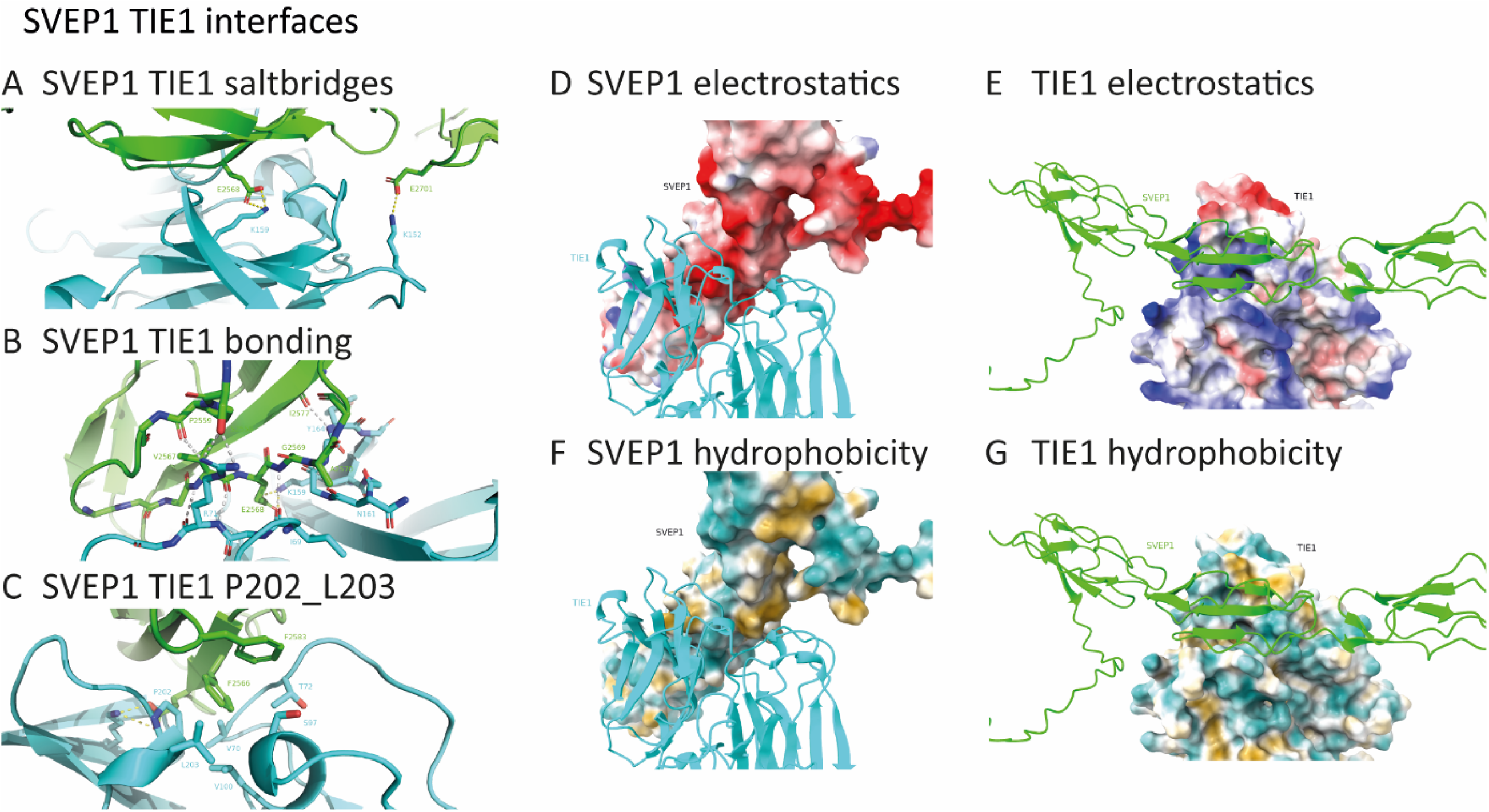
SVEP1 TIE1 interfaces. (**A**) Salt bridges E2568-K159 and E2701-K152 between SVEP1 CCP20 (green) and D1/D2 of TIE1 ectodomain (cyan). Electrostatic bonds are highlighted in yellow.. (**B**) Hydrogen bond network of SVEP1 CCP20 (green) and D1/D2 of TIE1 ectodomain (cyan). H-bonds are highlighted in white. (**C**) F2566 of SVEP1 (green) fits in the hydrophobic pocket of TIE1 (cyan) partially formed by P202 and L203. (**D** and **E**) Surface electrostatic potential representation of SVEP1 (green) bound to TIE1 (cyan) detailing that the mainly acidic interaction surface of SVEP1 (**D**) is opposed by a basic interaction surface of TIE1 (**E**). (**F** and **G**) Hydrophobicity surface representations of SVEP1 (green) bound to TIE1 (cyan). The hydrophobic side chains of F2566 and F2583 fit into a hydrophobic cavity of TIE1.

**Supplementary figure 1.3:**
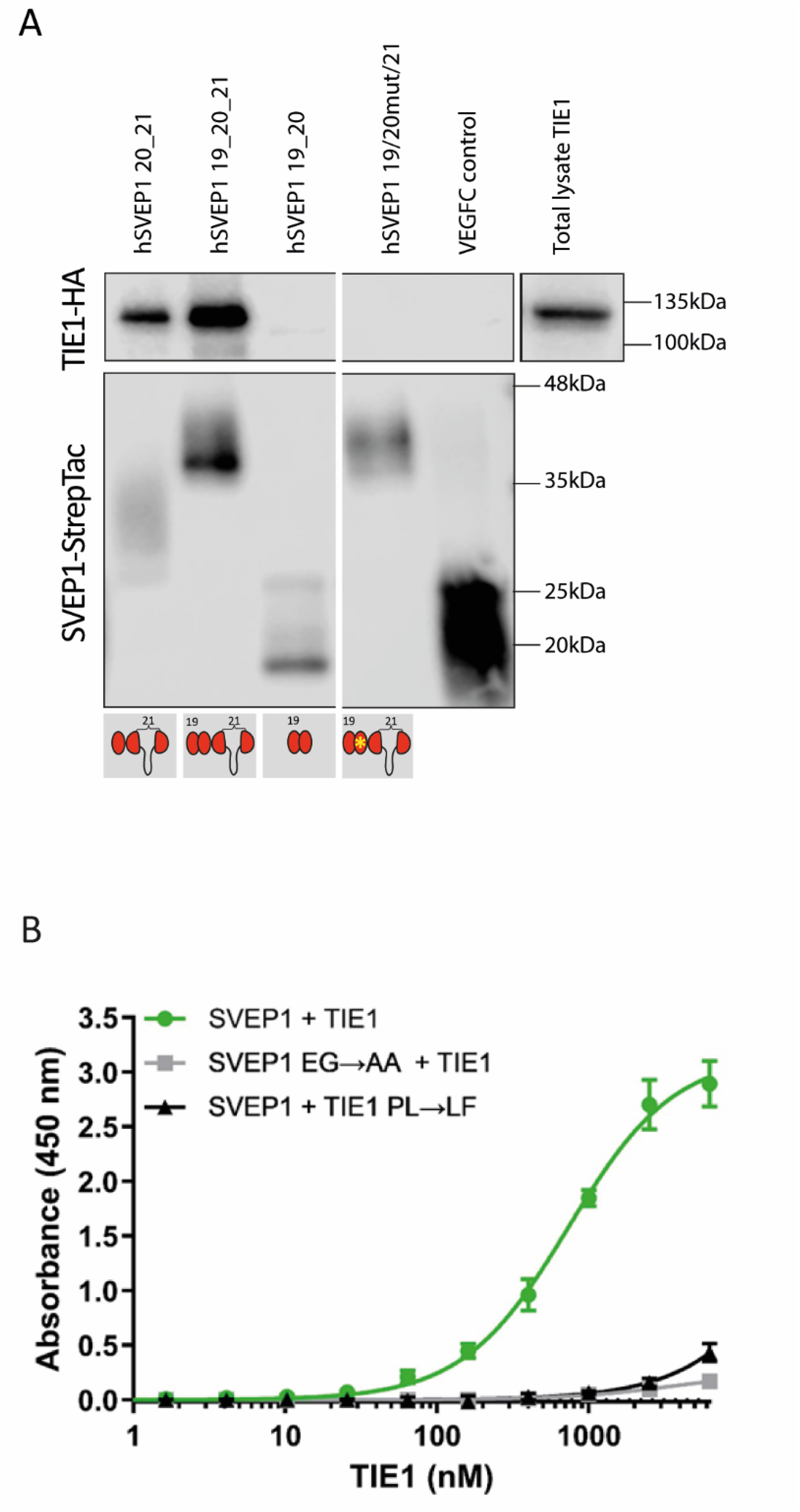
(**A**) Different versions of human SVEP1-StrepTac (spanning from CCP15 to the C-Term) were immunoprecipitated and associated TIE1 was detected via Western blot analysis. TIE1 bound to CCP19-21 as well as CCP20-21, but not the CCP19-21 mutant containing the E2568A-G2569A mutant or CCP19-20. (**B**) Binding of SVEP1 (wt or E2568A-G2569A) to TIE1 (wt or P202L L203F) was analyzed by ELISA. Both mutations, either in TIE1 or in SVEP1 inhibit binding between TIE1 and SVEP1.

**Supplementary figure 2.1:**
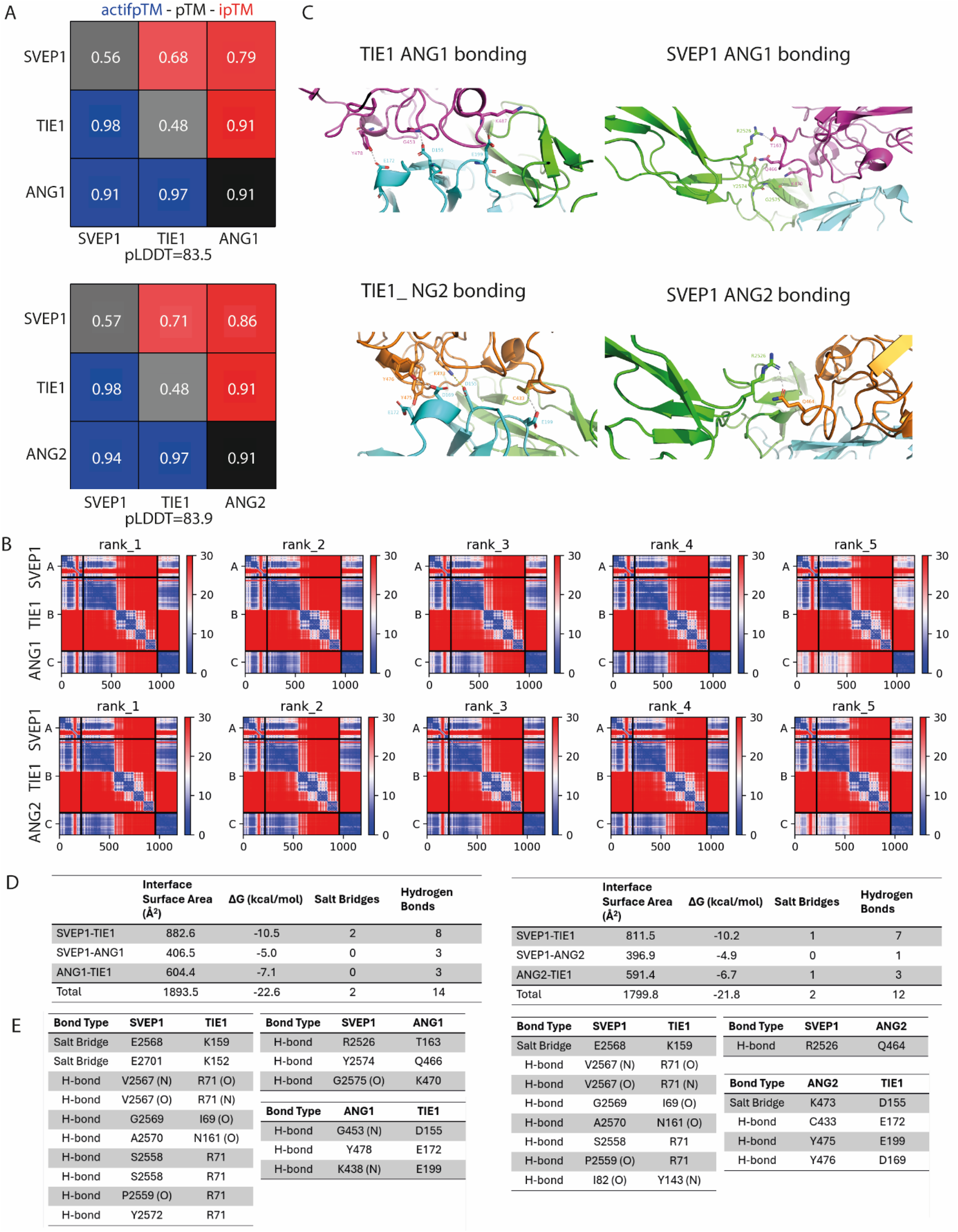
(**A**) AlphaFold 2.3 multimer output scores and (**B**) PAE plots of SVEP1-TIE1-ANG1/2 complexes. A is SVEP1 CCP19-21, B is TIE1 ecotdomain, C is the ANG1/2 fibrinogen like domain. (**C**) Bonding residues of each interface (**D**) and (**E**) PDBePISA analysis of interfaces of SVEP1-TIE1-ANG1/2 top ranked models. The conformation of the TIE1-ANG1/2 heteromers in the SVEP1-bound pose resembles the solved structures of TIE2 bound to either ANG1 or ANG2. However, the weaker binding of TIE2 to CCP19-21 of SVEP1 fails to nucleate the formation of the TIE2 ternary complex with SVEP1.

**Supplementary figure 2.2:**
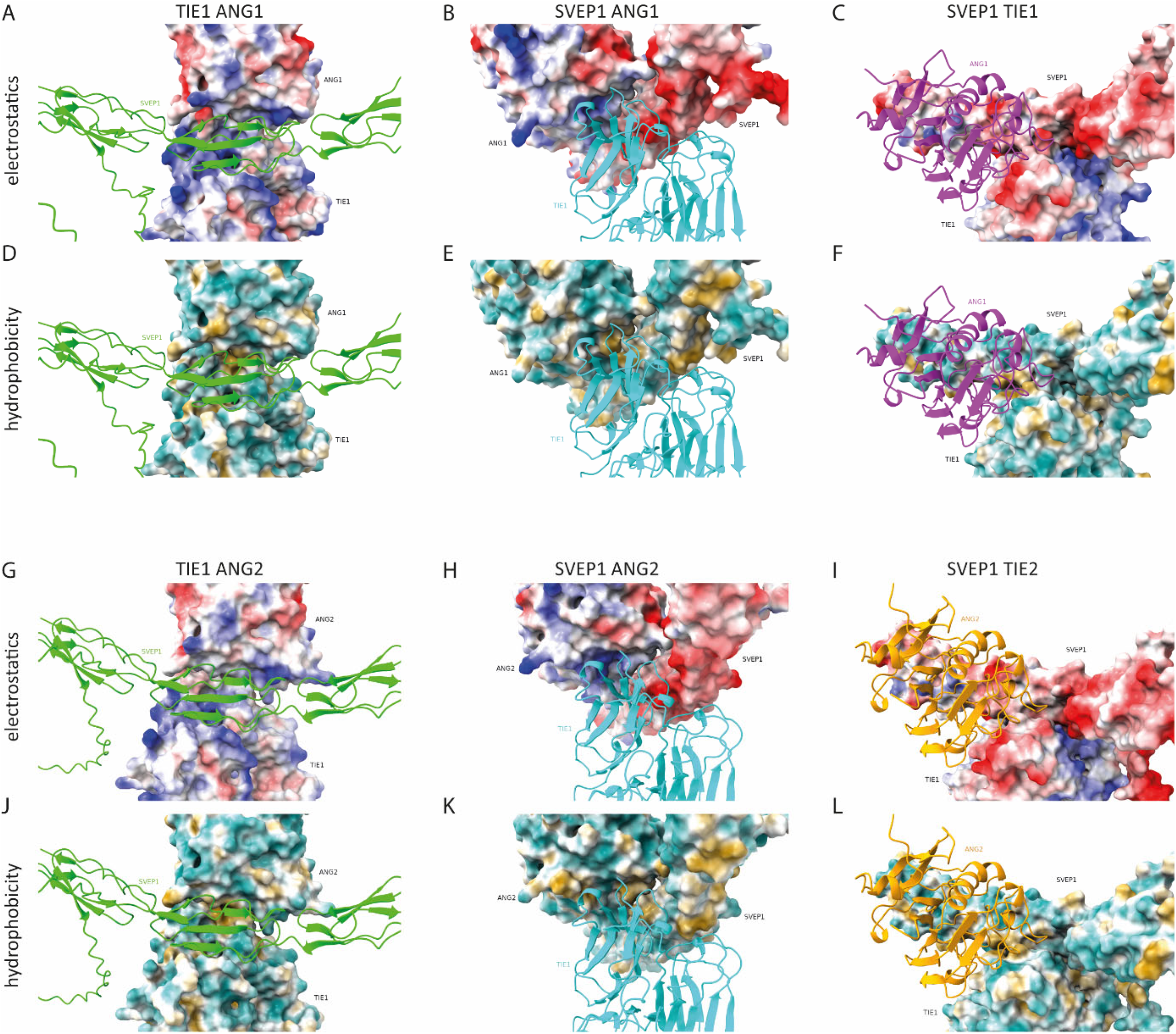
(A,B,C) Surface electrostatic potential of the interaction surfaces present in the SVEP1-TIE1-ANG1 complex demonstrating complementary charge distributions. (D,E,F) Surface hydrophobicity of the interaction surfaces present in the SVEP1-TIE1-ANG1 complex. (G,H,I) Surface electrostatic potential of the interaction surfaces present in the SVEP1-TIE1-ANG2 complex. (J,K,L) Surface hydrophobicity of the interaction surfaces present in the SVEP1-TIE1-ANG2 complex. All electrostatic potentials and hydrophobicity surfaces were generated and visualized with UCSF Chimera 1.19.

**Supplementary figure 2.3:**
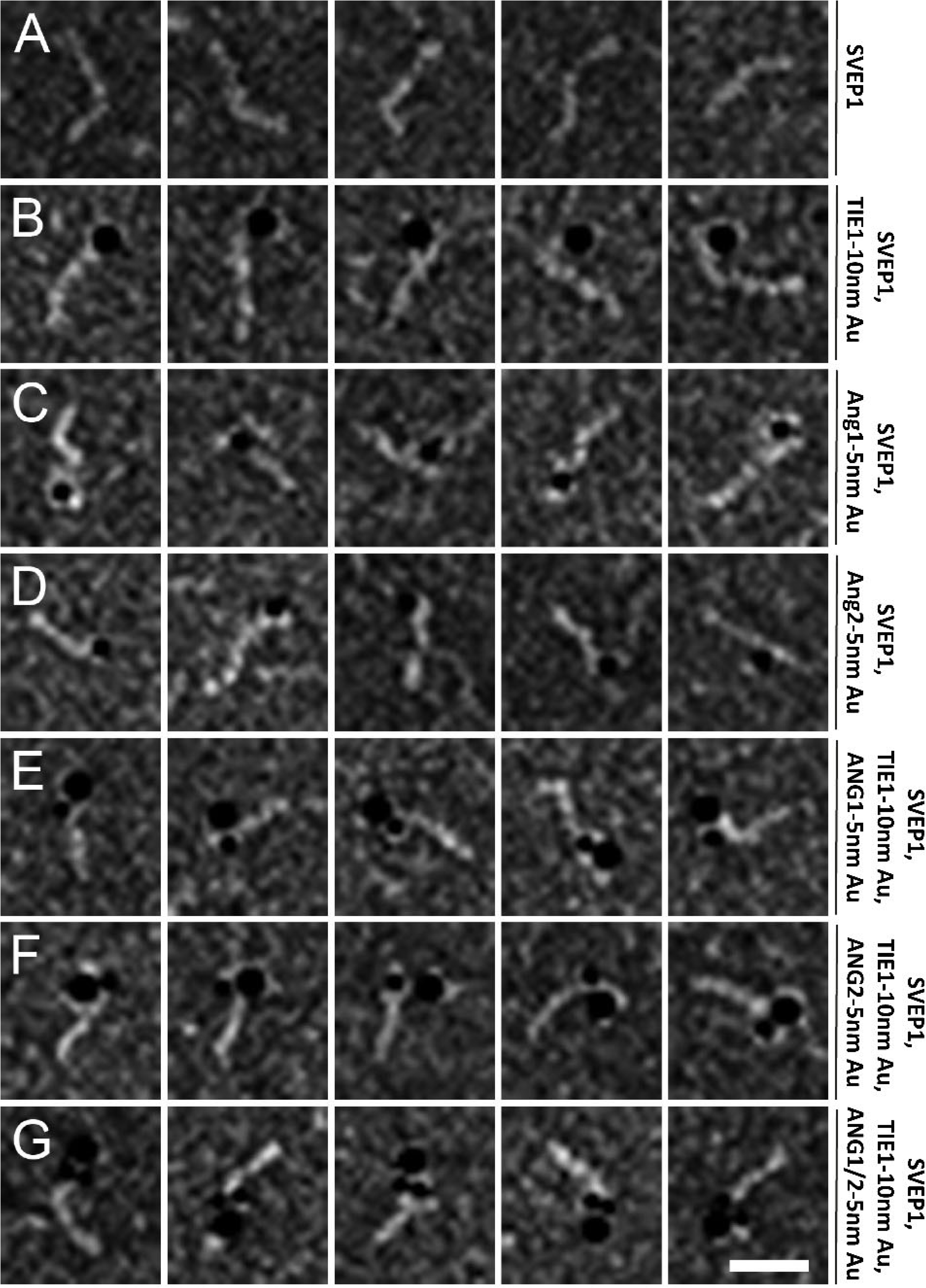
ANG1 and ANG2 bind in close proximity to the site where TIE1 and SVEP1 interact. Negative staining of human SVEP1 binding to TIE1 with and without the presence of ANG1 or ANG2. (**A**) hSVEP 150 kDa. (**B**) hSVEP-hTIE-10 nm Au. (**C**) hSVEP-ANG1-5nm Au, (**D**) hSVEP-ANG2-5nm Au. (**E**) hSVEP-hTIE-10 nm Au-ANG1-5 nm Au. (**F**) hSVEP-hTIE-10 nm Au-ANG2-5 nm Au. (**G**) hSVEP-hTIE-10 nm Au-ANG1/2-5 nm Au. Parts of panels A and B (control, SVEP1-TIE1-10 nm Au) have also been shown in main figure 1. Scale bar: 25 nm.

**Supplementary Figure 4.1:**
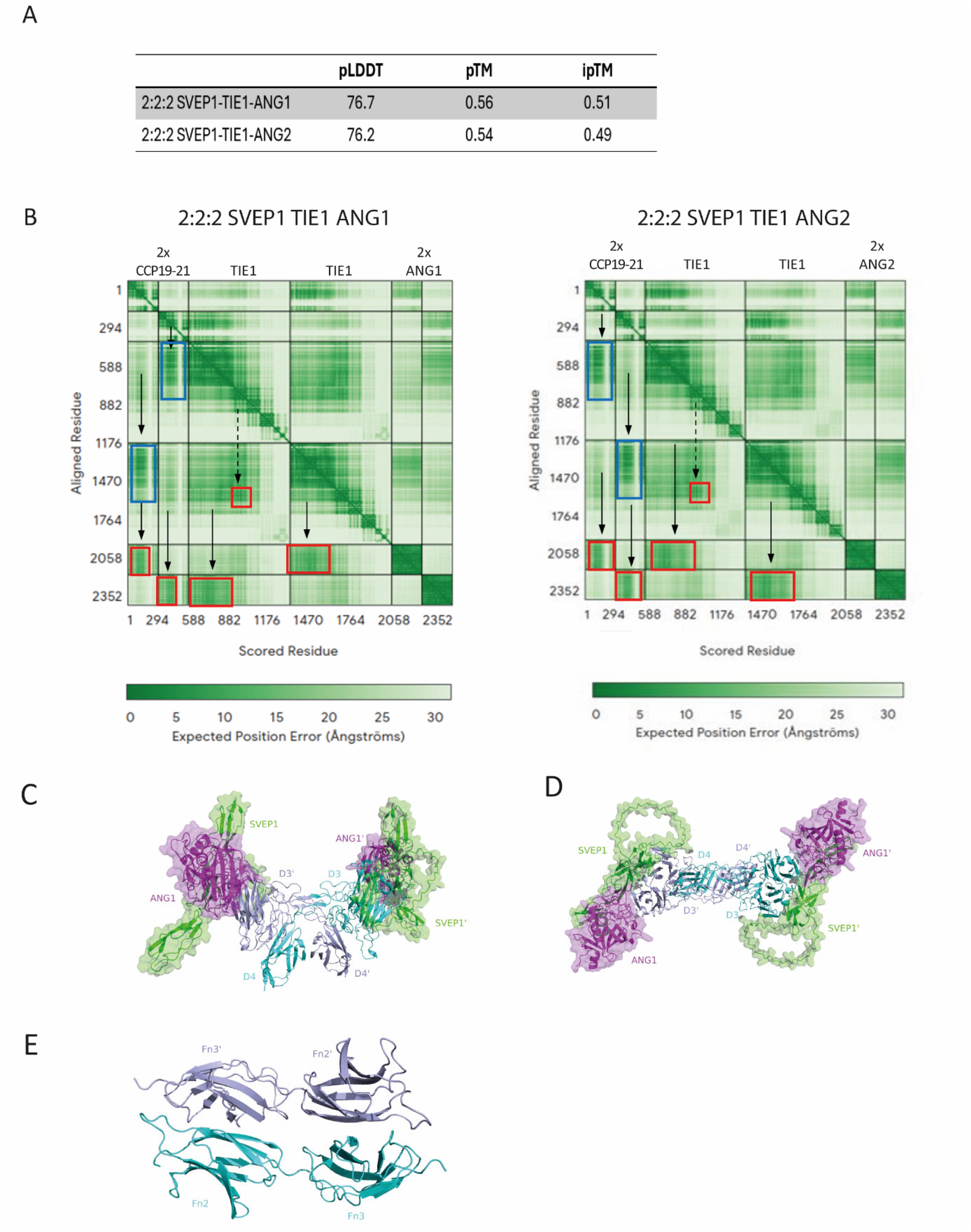
(A) AF3 output metrics (B) PAE plots demonstrating confident predictions of the complexes shown in Figure 4. (**C**) and (**D**) The side (**C**) and top (**D**) view of the model hexamer shows the side-by-side, antiparallel packing of TIE1 D3 domains that produces the domain-swapping of D4 Ig domains. **E**) Top view of the Fn2 – Fn3 domains.

## References

Aleström, Peter, Livia D’Angelo, Paul J. Midtlyng, Daniel F. Schorderet, Stefan Schulte-Merker, Frederic Sohm, and Susan Warner. 2020. “Zebrafish: Housing and Husbandry Recommendations.” Laboratory Animals 54 (3): 213–24. 10.1177/0023677219869037.

Baschong, W, and NG Wrigley. 1990. “Small colloidal gold conjugated to Fab fragments or to immunoglobulin G as high-resolution labels for electron microscopy: A technical overview.” Journal of electron microscopy technique 14 (4):313–323. doi:10.1002/jemt.1060140405.

Bos, Frank L., Maresa Caunt, Josi Peterson-Maduro, Lara Planas-Paz, Joe Kowalski, Terhi Karpanen, Andreas Van Impel, et al. 2011. “CCBE1 Is Essential for Mammalian Lymphatic Vascular Development and Enhances the Lymphangiogenic Effect of Vascular Endothelial Growth Factor-c in Vivo.” Circulation Research 109 (5): 486–91. 10.1161/CIRCRESAHA.111.250738.

Brouillard, Pascal, Aino Murtomäki, Veli Matti Leppänen, Marko Hyytiäinen, Sandrine Mestre, Lucas Potier, Laurence M. Boon, et al. 2024. “Loss-of-Function Mutations of the TIE1 Receptor Tyrosine Kinase Cause Late-Onset Primary Lymphedema.” The Journal of Clinical Investigation 134 (14). 10.1172/JCI173586.

Daly, Christopher, Vivian Wong, Elena Burova, Yi Wei, Stephanie Zabski, Jennifer Griffiths, Ka Man Lai, et al. 2004. “Angiopoietin-1 Modulates Endothelial Cell Function and Gene Expression via the Transcription Factor FKHR (FOXO1).” Genes & Development 18 (9): 1060–71. 10.1101/GAD.1189704.

Davis, Samuel, Thomas H. Aldrich, Pamela F. Jones, Ann Acheson, Debra L. Compton, Vivek Jain, Terence E. Ryan, et al. 1996. “Isolation of Angiopoietin-1, a Ligand for the TIE2 Receptor, by Secretion-Trap Expression Cloning.” Cell 87 (7): 1161–69. 10.1016/S0092-8674(00)81812-7.

Dumont, Daniel J., Gérard J. Gradwohl, Guo Hua Fong, Robert Auerbach, and Martin L. Breitman. 1993. “The Endothelial-Specific Receptor Tyrosine Kinase, Tek, Is a Member of a New Subfamily of Receptors.” Oncogene 8 (5): 1293–1301. https://europepmc.org/article/med/8386827.

Eble, Johannes A. 2018. “Titration ELISA as a Method to Determine the Dissociation Constant of Receptor Ligand Interaction.” J Vis Exp (132). doi: 10.3791/57334.

Gjini, Evisa, Liesbeth H Hekking, Axel Küchler, Pipsa Saharinen, Erno Wienholds, Jan-Andries Post, Kari Alitalo, and Stefan Schulte-Merker. 2011. “Zebrafish Tie-2 shares a redundant role with Tie-1 in heart development and regulates vessel integrity.” Disease models & mechanisms 4 (1):57–66. doi: 10.1242/dmm.005033.

Guen, Ludovic Le, Terhi Karpanen, Dörte Schulte, Nicole C. Harris, Katarzyna Koltowska, Guy Roukens, Neil I. Bower, et al. 2014. “Ccbe1 Regulates Vegfc-Mediated Induction of Vegfr3 Signaling during Embryonic Lymphangiogenesis.” Development 141 (6): 1239–49. 10.1242/DEV.100495.

Hansen, Tania M., Harprit Singh, Tariq A. Tahir, and Nicholas P.J. Brindle. 2010. “Effects of Angiopoietins-1 and -2 on the Receptor Tyrosine Kinase Tie2 Are Differentially Regulated at the Endothelial Cell Surface.” Cellular Signalling 22 (3): 527–32. 10.1016/J.CELLSIG.2009.11.007.

Hogan, Benjamin M., Robert Herpers, Merlijn Witte, Hanna Heloterä, Kari Alitalo, Henricus J. Duckers, and Stefan Schulte-Merker. 2009. “Vegfc/Flt4 Signalling Is Suppressed by Dll4 in Developing Zebrafish Intersegmental Arteries.” Development 136 (23): 4001–9. 10.1242/dev.039990.

Hußmann, Melina, Dörte Schulte, Sarah Weischer, Claudia Carlantoni, Hiroyuki Nakajima, Naoki Mochizuki, Didier Y.R. Stainier, Thomas Zobel, Manuel Koch, and Stefan Schulte-Merker. 2023. “Svep1 Is a Binding Ligand of Tie1 and Affects Specific Aspects of Facial Lymphatic Development in a Vegfc-Independent Manner.” ELife 12 (April): e82969. 10.7554/ELIFE.82969.

Impel, Andreas van, Zhonghua Zhao, Dorien M.A. Hermkens, M. Guy Roukens, Johanna C. Fischer, Josi Peterson-Maduro, Henricus Duckers, Elke A. Ober, Philip W. Ingham, and Stefan Schulte-Merker. 2014. “Divergence of Zebrafish and Mouse Lymphatic Cell Fate Specification Pathways.” Development (Cambridge) 141 (6): 1228–38. 10.1242/dev.105031.

Jeltsch, Michael, Sawan Kumar Jha, Denis Tvorogov, Andrey Anisimov, Veli Matti Leppänen, Tanja Holopainen, Riikka Kivelä, Sagrario Ortega, Terhi Kärpanen, and Kari Alitalo. 2014. “CCBE1 Enhances Lymphangiogenesis via a Disintegrin and Metalloprotease with Thrombospondin Motifs-3-Mediated Vascular Endothelial Growth Factor-C Activation.” Circulation 129 (19): 1962–71. 10.1161/CIRCULATIONAHA.113.002779.

Jiang, Zhen, Claudia Carlantoni, Srinivas Allanki, Ingo Ebersberger, and Didier Y.R. Stainier. 2020. “Tek (Tie2) Is Not Required for Cardiovascular Development in Zebrafish.” Development (Cambridge, England) 147 (19). 10.1242/DEV.193029.

Karpanen, Terhi, Yvonne Padberg, Serge A. Van De Pavert, Cathrin Dierkes, Nanami Morooka, Josi Peterson-Maduro, Glenn Van De Hoek, et al. 2017. “An Evolutionarily Conserved Role for Polydom/Svep1 during Lymphatic Vessel Formation.” Circulation Research 120 (8). 10.1161/CIRCRESAHA.116.308813.

Kim, Hak-Zoo, Keehoon Jung, Ho Min Kim, Yifan Cheng, and Gou Young Koh. 2009. “A designed angiopoietin-2 variant, pentameric COMP-Ang2, strongly activates Tie2 receptor and stimulates angiogenesis.” Biochimica et Biophysica Acta (BBA)-Molecular Cell Research 1793 (5):772–780. doi: 10.1016/j.bbamcr.2009.01.018.

Kim, Injune, Hwan Gyu Kim, June-No So, Joo Heon Kim, Hee Jin Kwak, and Gou Young Koh. 2000. “Angiopoietin-1 regulates endothelial cell survival through the phosphatidylinositol 3ʹ-kinase/Akt signal transduction pathway.” Circulation research 86 (1):24–29. doi: 10.1161/01.res.86.1.24.

Kim, Kyung-Tae, Han-Ho Choi, Michel O Steinmetz, Bohumil Maco, Richard A Kammerer, So Young Ahn, Hak-Zoo Kim, Gyun Min Lee, and Gou Young Koh. 2005. “Oligomerization and multimerization are critical for angiopoietin-1 to bind and phosphorylate Tie2.” Journal of Biological Chemistry 280 (20):20126–20131. doi: 10.1074/jbc.M500292200.

Korhonen, Emilia A., Anita Lampinen, Hemant Giri, Andrey Anisimov, Minah Kim, Breanna Allen, Shentong Fang, et al. 2016. “Tie1 Controls Angiopoietin Function in Vascular Remodeling and Inflammation.” The Journal of Clinical Investigation 126 (9): 3495. 10.1172/JCI84923.

Korhonen, Emilia A., Aino Murtomaki, Sawan Kumar Jha, Andrey Anisimov, Anne Pink, Yan Zhang, Simon Stritt, et al. 2022. “Lymphangiogenesis Requires Ang2/Tie/PI3K Signaling for VEGFR3 Cell-Surface Expression.” The Journal of Clinical Investigation 132 (15). 10.1172/JCI155478.

Krissinel, Evgeny and Henrick, Kim. 2007. “Inference of macromolecular assemblies from crystalline state.” J Mol Biol 372 (3):774–97. doi: 10.1016/j.jmb.2007.05.022.

Kumar, Sanjay Sunil, Katharina Uphoff, Sophie Hötte, Verena Prokosch, Stefan Schulte-Merker, and Dörte Schulte-Ostermann. 2026. “A cellular and molecular perspective on organotypic lymphatic (dys) function.” Seminars in Cell & Developmental Biology. 10.1016/j.semcdb.2025.103665

Leppänen, Veli Matti, Pipsa Saharinen, and Kari Alitalo. 2017. “Structural Basis of Tie2 Activation and Tie2/Tie1 Heterodimerization.” Proceedings of the National Academy of Sciences of the United States of America 114 (17): 4376–81. 10.1073/PNAS.1616166114.

Macdonald, Philip R., Pavlos Progias, Barbara Ciani, Sanjai Patel, Ulrike Mayer, Michel O. Steinmetz, and Richard A. Kammerer. 2006. “Structure of the Extracellular Domain of Tie Receptor Tyrosine Kinases and Localization of the Angiopoietin-Binding Epitope.” Journal of Biological Chemistry 281 (38): 28408–14. 10.1074/JBC.M605219200.

Maisonpierre, Peter C., Chitra Suri, Pamela F. Jones, Sona Bartunkova, Stanley J. Wiegand, Czeslaw Radziejewski, Debra Compton, et al. 1997. “Angiopoietin-2, a Natural Antagonist for Tie2 That Disrupts in Vivo Angiogenesis.” Science 277 (5322): 55–60. 10.1126/SCIENCE.277.5322.55

Mäkinen, Taija, Laurence M. Boon, Miikka Vikkula, and Kari Alitalo. 2021. “Lymphatic Malformations: Genetics, Mechanisms and Therapeutic Strategies.” Circulation Research 129 (1): 136–54. 10.1161/CIRCRESAHA.121.318142

Marron, Marie B., David P. Hughes, Michael D. Edge, Cheryl L. Forder, and Nicholas P.J. Brindle. 2000. “Evidence for Heterotypic Interaction between the Receptor Tyrosine Kinases Tie-1 and Tie-2.” Journal of Biological Chemistry 275 (50): 39741–46. 10.1074/jbc.M007189200.

Meier, Markus, Monika Gupta, Serife Akgül, Matthew McDougall, Thomas Imhof, Denise Nikodemus, Raphael Reuten, Aniel Moya-Torres, Vu To, Fraser Ferens, Fabian Heide, Gay Pauline Padilla-Meier, Philipp Kukura, Wenming Huang, Birgit Gerisch, Matthias Mörgelin, Kate Poole, Adam Antebi, Manuel Koch & Jörg Stetefeld. 2023. “The dynamic nature of netrin-1 and the structural basis for glycosaminoglycan fragment-induced filament formation.” Nat Commun 14 (1):1226. doi: 10.1038/s41467-023-36692-w.

Moore, Jason O, Mark A Lemmon, and Kathryn M Ferguson. 2017. “Dimerization of Tie2 mediated by its membrane-proximal FNIII domains.” Proceedings of the National Academy of Sciences 114 (17):4382–4387. doi: 10.1073/pnas.1617800114.

Morooka, Nanami, Sugiko Futaki, Ryoko Sato-Nishiuchi, Masafumi Nishino, Yuta Totani, Chisei Shimono, Itsuko Nakano, Hiroyuki Nakajima, Naoki Mochizuki, and Kiyotoshi Sekiguchi. 2017. “Polydom Is an Extracellular Matrix Protein Involved in Lymphatic Vessel Remodeling.” Circulation Research 120 (8): 1276–88. 10.1161/CIRCRESAHA.116.308825.

Morooka, Nanami, Ning Gui, Koji Ando, Keisuke Sako, Moe Fukumoto, Urara Hasegawa, Melina Hußmann, Stefan Schulte-Merker, Naoki Mochizuki, and Hiroyuki Nakajima. 2024. “Angpt1 Binding to Tie1 Regulates the Signaling Required for Lymphatic Vessel Development in Zebrafish.” Development (Cambridge) 151 (10). 10.1242/DEV.202269.

Partanen, Juha, Elina Armstrong, Tomi P. Mäkelä, Jaana Korhonen, Minna Sandberg, Risto Renkonen, Sakari Knuutila, Kay Huebner, and Kari Alitalo. 1992. “A Novel Endothelial Cell Surface Receptor Tyrosine Kinase with Extracellular Epidermal Growth Factor Homology Domains.” Molecular and Cellular Biology 12 (4): 1698–1707. 10.1128/MCB.12.4.1698-1707.1992.

Petrova, Tatiana V., and Gou Young Koh. 2018. “Organ-Specific Lymphatic Vasculature: From Development to Pathophysiology.” Journal of Experimental Medicine 215 (1): 35–49. 10.1084/jem.20171868.

Roukens, M. Guy, Josi Peterson-Maduro, Yvonne Padberg, Michael Jeltsch, Veli Matti Leppänen, Frank L. Bos, Kari Alitalo, Stefan Schulte-Merker, and Dörte Schulte. 2015. “Functional Dissection of the CCBE1 Protein: A Crucial Requirement for the Collagen Repeat Domain.” Circulation Research 116 (10): 1660–69. 10.1161/CIRCRESAHA.116.304949.

Saharinen, Pipsa, Lauri Eklund, and Kari Alitalo. 2017. “Therapeutic Targeting of the Angiopoietin–TIE Pathway.” Nature Reviews Drug Discovery 2017 16:9 16 (9): 635–61. 10.1038/nrd.2016.278.

Saharinen, Pipsa, Katja Kerkelä, Niklas Ekman, Marie Marron, Nicholas Brindle, Gyun Min Lee, Hellmut Augustin, Gou Young Koh, and Kari Alitalo. 2005. “Multiple Angiopoietin Recombinant Proteins Activate the Tie1 Receptor Tyrosine Kinase and Promote Its Interaction with Tie2.” The Journal of Cell Biology 169 (2): 239. 10.1083/JCB.200411105.

Sato-Nishiuchi, Ryoko, Masamichi Doiguchi, Nanami Morooka, and Kiyotoshi Sekiguchi. 2023. “Polydom/SVEP1 Binds to Tie1 and Promotes Migration of Lymphatic Endothelial Cells.” Journal of Cell Biology 222 (9). 10.1083/JCB.202208047.

Sato-Nishiuchi, Ryoko, Itsuko Nakano, Akio Ozawa, Yuya Sato, Makiko Takeichi, Daiji Kiyozumi, Kiyoshi Yamazaki, Teruo Yasunaga, Sugiko Futaki, and Kiyotoshi Sekiguchi. 2012. “Polydom/SVEP1 Is a Ligand for Integrin Α9β1.” Journal of Biological Chemistry 287 (30): 25615–30. 10.1074/jbc.M112.355016.

Savant, Soniya, Silvia La Porta, Annika Budnik, Katrin Busch, Junhao Hu, Nathalie Tisch, Claudia Korn, et al. 2015. “The Orphan Receptor Tie1 Controls Angiogenesis and Vascular Remodeling by Differentially Regulating Tie2 in Tip and Stalk Cells.” Cell Reports 12 (11): 1761. 10.1016/J.CELREP.2015.08.024.

Seegar, Tom C.M., Becca Eller, Dorothea Tzvetkova-Robev, Momchil V. Kolev, Scott C. Henderson, Dimitar B. Nikolov, and William A. Barton. 2010. “Tie1-Tie2 Interactions Mediate Functional Differences between Angiopoietin Ligands.” Molecular Cell 37 (5): 643–55. 10.1016/j.molcel.2010.02.007.

Shen, Bin, Zhi Shang, Bo Wang, Luqing Zhang, Fei Zhou, Taotao Li, Man Chu, et al. 2014. “Genetic Dissection of Tie Pathway in Mouse Lymphatic Maturation and Valve Development.” Arteriosclerosis, Thrombosis, and Vascular Biology 34 (6): 1221–30. 10.1161/ATVBAHA.113.302923.

Souma, Tomokazu, Benjamin R. Thomson, Stefan Heinen, Isabel Anna Carota, Shinji Yamaguchi, Tuncer Onay, Pan Liu, et al. 2018. “Context-Dependent Functions of Angiopoietin 2 Are Determined by the Endothelial Phosphatase VEPTP.” Proceedings of the National Academy of Sciences of the United States of America 115 (6): 1298–1303. 10.1073/PNAS.1714446115.

Wang, Guangxia, Lars Muhl, Yvonne Padberg, Laura Dupont, Josi Peterson-Maduro, Martin Stehling, Ferdinand le Noble, et al. 2020. “Specific Fibroblast Subpopulations and Neuronal Structures Provide Local Sources of Vegfc-Processing Components during Zebrafish Lymphangiogenesis.” Nature Communications 11 (1). 10.1038/s41467-020-16552-7.

Young, Terri L., Kristina N. Whisenhunt, Jing Jin, Sarah M. LaMartina, Sean M. Martin, Tomokazu Souma, Vachiranee Limviphuvadh, et al. 2020. “SVEP1 as a Genetic Modifier of TEK-Related Primary Congenital Glaucoma.” Investigative Ophthalmology and Visual Science 61 (12). 10.1167/IOVS.61.12.6.

Yuan, Hai Tao, Shivalingappa Venkatesha, Barden Chan, Urban Deutsch, Tadanori Mammoto, Vikas P. Sukhatme, Adrian S. Woolf, and S. Ananth Karumanchi. 2007. “Activation of the Orphan Endothelial Receptor Tie1 Modifies Tie2-mediated Intracellular Signaling and Cell Survival.” The FASEB Journal 21 (12): 3171–83. 10.1096/fj.07-8487com.

